# Adaptive tracking contributes to the invasion success of a global pest insect

**DOI:** 10.64898/2026.02.26.708249

**Authors:** Eleanor A. McCabe, Mathieu Gautier, Kit A. Eller, Miles O. Garvin, Andrew R. McCracken, Corey Forman, Michael Darrington, Mark Garcia, Sebastian Redondo, Alan O. Bergland, Alyssa Bangerter, Katie E. Lotterhos, Joaquin C. B. Nunez, Nicholas M. Teets

## Abstract

Invasive species are a threat to biodiversity, natural resources, and the economy. Preventing future invasions requires the ability to predict which species will be successful invaders. Invading a new region often requires coping with novel environmental conditions, yet how invading populations adapt to environmental conditions encountered in their introduced range is not always clear. One process that may facilitate invasion in temperate environments is seasonal adaptive tracking, a rapid evolutionary process in which allele frequencies and phenotypes shift in response to seasonal environmental fluctuations, promoting the long-term maintenance of polymorphisms. Here, we test the hypothesis that adaptive tracking is a key contributor to the invasion success of the globally invasive fly *Drosophila suzukii*. We demonstrate that adaptive tracking results in predictable, rapid evolution of a key thermal tolerance trait (critical thermal minimum) essential for coping with climate variability. We also detect adaptive tracking in several genomic regions enriched for seasonally fluctuating alleles, and these alleles map to loci implicated in diverse traits, including thermal tolerance, pesticide resistance, and olfactory behavior. Finally, we find that loci undergoing adaptive tracking are significantly enriched for alleles associated with invasion in North American populations, suggesting that rapid evolution of seasonal traits may contribute to the success of this invasive species.

**Significance Statement:** Understanding how invasive species succeed in a new environment is important for forecasting and managing new invasions. Here we show that rapid evolution in response to changing seasons contributes to the success of the highly invasive pest *Drosophila suzukii*. In wild populations, cold tolerance rapidly evolves on yearly timescales, and evolution of this important seasonal trait may help *D. suzukii* adapt to novel climates. We also observe clear evidence of adaptive tracking throughout the genome, including genes related to several traits that may be important to seasonality. Further, many of the alleles undergoing adaptive tracking overlap with alleles known to be important for invasion, indicating that adaptive tracking is an important evolutionary mechanism contributing to the success of invasive species.

## Introduction

Invasive species are an increasingly growing problem and present a significant threat to biodiversity, agriculture, and the economy [1–3]. In North America alone, invasives cost the economy approximately $30 billion per year, and nearly half that cost is experienced by the agriculture sector [4]. The number of introduced species far exceeds the number that successfully establish and spread [5, 6], and identifying characteristics of successful invaders can inform management decisions to minimize the impact of future invasions.

Invasive species face novel abiotic and biotic challenges in their expanded range. While pre-existing life history traits that promote successful invasion are well-characterized [7–9], it is becoming increasingly apparent that rapid evolution is important during invasion [10]. Specifically, the evolutionary dynamics of invasive species are shaped by multiple processes, including founder effects, repeated introductions, and natural selection. Adaptive tracking, another evolutionary process that can facilitate rapid adaptation, may provide an additional way for invasive species to adapt to seasonally variable habitats. Adaptive tracking occurs when fluctuating environmental conditions cause selection on standing genetic variation, which drives predictable fluctuations in allele frequencies that confer context dependent fitness advantages [11]. Thus, adaptive tracking can preserve functional polymorphism [12] and allow populations to maximize fitness across changing environments. However, whether the genetic mechanisms that underlie adaptive tracking are the same as those that underlie invasion success has not been empirically tested. In this study we ask the questions: (i) does adaptive tracking contribute to seasonality in a recently invaded species, and (ii) do adaptive tracking and invasiveness share a genetic basis?

Adaptive tracking has been demonstrated in several cosmopolitan multivoltine species, which undergo predictable seasonal shifts in allele frequency that correlate with seasonal changes in temperature [13–17]. In *Drosophila melanogaster,* these changes in allele frequencies correspond with changes in adaptive phenotypes like reproductive traits, stress tolerance, immunity, and insecticide resistance [17–19]. However, few species have been assessed for the capacity to undergo adaptive tracking [20], and even among these, adaptive tracking is not universal [21]. Further, much of the phenotypic evidence for adaptive tracking was observed in caged populations [17, 18, 22, 23], and assessing the ecological relevance of this process requires studies in wild populations where gene flow and other evolutionary drivers are unrestricted. Nonetheless, by allowing for real-time optimization of phenotypes in fluctuating environments, adaptive tracking may be an important part of the toolkit for invasive species.

*Drosophila suzukii* (Matsumura, 1931) provides an excellent system for testing the hypothesis that adaptive tracking is a key contributor of invasion success. As one of the most invasive insects in recent history, this species colonized nearly all of the continental US and much of Canada over an ∼8-year period [24]. Females lay eggs directly into fruit still ripening on plants, and feeding damage by larvae causes millions of dollars of yield loss and management costs each year [25]. Its current range spans a broad diversity of continental climates, suggesting an ability to rapidly adapt to novel climatic conditions. *D. suzukii* has the necessary ingredients for seasonal adaptive tracking, as it can complete ∼10 generations per year in temperate latitudes [26]. Furthermore, the population genetic characteristics of *D. suzukii* have been assessed on a global scale [27, 28], including identification of alleles important for invasion success [28, 29]. Thus, the life history traits, agronomic impacts, and genetic resources of *D. suzukii* are ideal for empirically determining the role of adaptive tracking during its worldwide invasion.

In this study we tested whether adaptive tracking is a factor in the invasion success of *D. suzukii.* We first established the three components needed for a species to undergo adaptive tracking: 1) a stable, year-round resident population exposed to temporal environmental variation, 2) phenotypic changes in response to these fluctuating environments, and 3) consistent temporal changes in allele frequency at functional polymorphisms. Specifically, we determined seasonal changes in allele frequencies and an ecologically important thermal tolerance trait, critical thermal minimum (CT_min_). We repeatedly sampled a wild population over four consecutive years, genotyping and phenotyping individuals at three time points per year to demonstrate that *D. suzukii* is undergoing seasonal adaptive tracking. We further show that genes under seasonal selection are enriched among those associated with invasion success, indicating that adaptive tracking shares mechanisms associated with the invasive spread and persistence of *D. suzukii*.

## Results and Discussion

### Seasonal and temporal demography of *D. suzukii*’s invasion history

We used pooled sequencing of global *D. suzukii* samples [28], including new samples generated here (i.e., 19 temporal pool-seq samples from two Kentucky sites spanning 2020 to 2023, and 4 temporal pool-seq samples from Charlottesville, Virginia spanning 2017-2018; **Tables S1-S3; Dataset S1**), to resolve population structure shaped by the species’ invasion history **Fig. 1A**). Global principal component analysis (PCA) showed discrete population clusters corresponding to native and invasive ranges. This pattern is consistent with the hypothesis of two parallel, yet independent, invasion routes [28, 30], eastward into the Americas and westward into Europe (**Fig. 1B**). Next, we leveraged the North American samples to test for signatures of spatial and temporal stability, a key prerequisite for adaptive tracking in seasonal environments. To assess population stability through time, we performed a principal component analysis (PCA) of genome-wide genetic variation. Within North America, patterns of population structure also appear to reflect the trajectory of invasion, with the first principal component (PC 1, 4.88%; **Fig. 1C**) primarily separating populations east and west of the Rocky Mountains, consistent with an introduction to the West Coast from the Asian native area and Hawaii (**Fig. 1D**; ANOVA; *F*_1,26_ = 147.9, *P* = 3.11×10^−12^) [31, 32]. These results, together with demographic modeling of Kentucky and Virginia populations (see ***supplemental text S1***), indicate that North American populations are spatially structured yet connected by ongoing gene flow, consistent with previous studies of *D. suzukii* genetic structure [27, 33].

**Figure 1.**
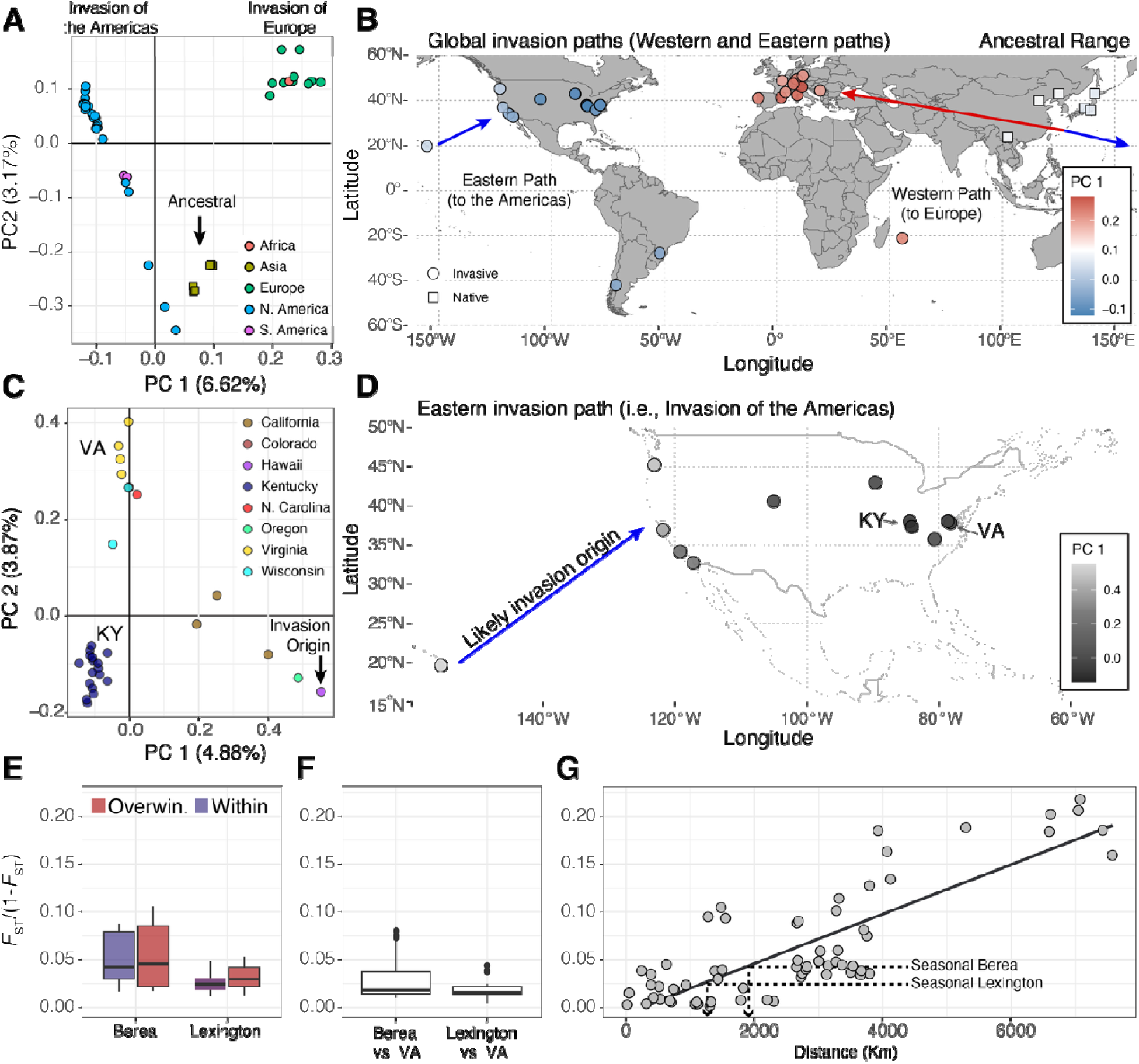
Spatial and Temporal Demography of *D. suzukii*’s Worldwide Invasions. **(A)** Principal component analysis (PCA) of worldwide SWD samples. The arrows represent the broad invasion routes for SWD. **(B)** PC 1 loadings of the worldwide analysis projected onto a world map based on the sampling coordinates of all samples. **(C)** Principal component analysis (PCA) of North American SWD samples. Here, the arrow represents the likely invasion origin. **(D)** PC 1 loadings of the North American analysis projected onto a world map based on the sampling coordinates of the samples**. (E)** Temporal *F*_ST_ estimates for seasonal KY samples. In this context, overwintering *F*_ST_ (i.e., “Overwin.”) represents the levels of genetic differentiation among samples collected from different years, and within year *F*_ST_ (i.e., “Within”) represents genetic differentiation among samples collected within the same growing season. Overwintering *F*_ST_ is a proxy for the severity of winter bottlenecks in temperate Drosophila [12]. **(F)** Spatial *F*_ST_ between our two sampling localities in Kentucky and our samples in Virginia**. (G)** Relationship between Haversine distance and FST across all North American samples. For panels E, F, and G, all *F*_ST_ values were linearized as *F*_ST_ /(1 − *F*_ST_).

Capitalizing on the temporal resolution of our Kentucky sampling, we quantified genetic differentiation (*F*_ST_) to assess patterns of population permanence across time and space. Temporal differentiation ranged from *F*_ST_ = 0.025 to 0.040 and remained consistent both within and across years (**Fig. 1E**). This pattern contrasts with other temperate drosophilids (i.e., *D. melanogaster*), where strong overwintering bottlenecks (∼90% population contraction) generate greater differentiation among years than within years and are a major driver of temporal genetic variation [12]. In *D. suzukii*, by contrast, the absence of elevated among-year differentiation suggests that overwintering bottlenecks may be considerably weaker (see ***Supplementary Text S2***).

To contextualize this differentiation, we compared temporal *F_ST_*within Kentucky with spatial differentiation across North American populations. Strikingly, differentiation between Kentucky and neighboring Virginia populations was lower (*F_ST_* = 0.023; **Fig. 1F**) than that among Kentucky samples collected at different times (*F*_ST_ ≥0.025). Within a single year, Kentucky populations can become as differentiated as populations separated by approximately ∼1,500 km (**Fig. 1G**, equivalent to ∼13° of latitude). While our demographic modeling suggests there is ongoing migration among neighboring populations (***supplemental text S1***), the patterns of spatial stability seen in the PCA (**Fig. 1C**; i.e., Kentucky samples clustered together irrespective of collection year), and in our demographic modeling (***supplemental text S1***), lead us to hypothesize that temporal genetic differentiation in *D. suzukii* may be generated, at least in part, by seasonal adaptive tracking operating within locally persistent populations [12, 13].

### Seasonal evolutionary change in CT_min_

To assess if adaptive tracking is occurring in *D. suzukii,* we conducted parallel phenotypic and genetic characterizations of *D. suzukii* sampled three times per year, approximately six weeks apart, for four consecutive years at two farms in central Kentucky (**Fig. 2A**). The first and third sampling intervals were approximately 5 generations apart based on degree day estimates. We allowed flies to emerge from infested fruit and quantified CT_min_ in both the parental (P) generation and in the F4 generation of isofemale lines derived from these collections. Flies from the P generation were also used for genetic analyses. CT_min_ measurements in the F4 generation were conducted by pooling equal numbers of flies from up to 20 isofemale lines from each collection (see *supplemental methods*; **Table S1**).

**Figure 2.**
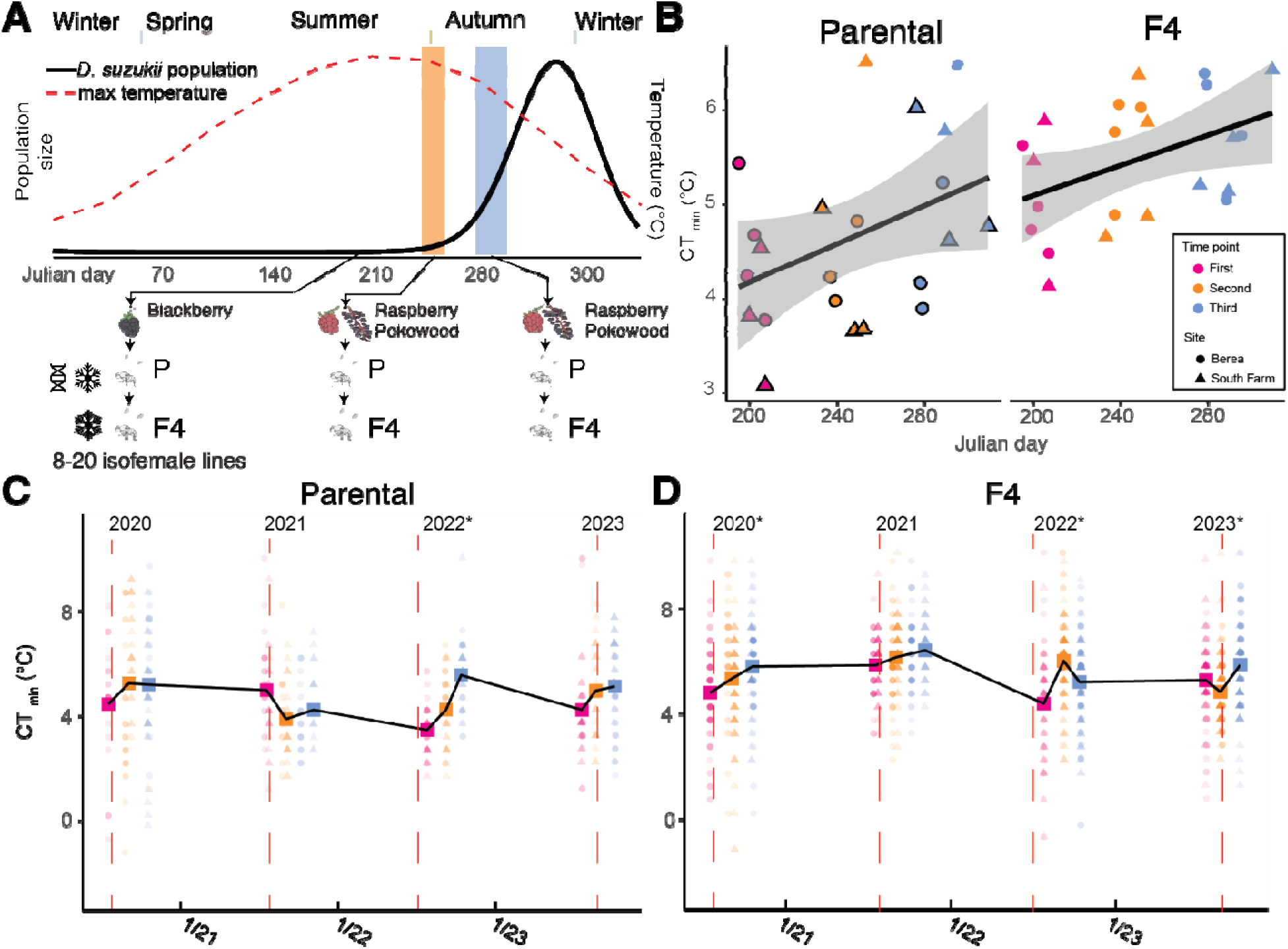
Evidence for adaptive tracking of CT_min_ in *D. suzukii*. **(A)** Schematic of sampling scheme. The range of sampling dates for time points between the different years is indicated with vertical rectangles, while the black line indicates predicted relative population size in central KY, as modeled by McCabe et al 2025 [26]. The red dashed line indicates average maximum temperature per month for central Kentucky. From each collection, we measured CT_min_ and genotyped the P generation and isofemale lines were propagated until the F4 generation, and phenotyped again. **(B)** Mean CT_min_ by Julian date. Points outlined in black were groups used for genotyping**. (C)** CT_min_ for each sampling point in the P generation. Each point represents a single fly, with circles representing Berea and triangles representing Lexington. Larger squares are combined means from both locations, based on n = 14-31 individuals. Dashed vertical red lines represent the hottest day of the year. Years marked with an asterisk (*) indicate a significant increase in CT_min_, based on Tukey’s HSD post hoc analysis of the sampling points within each year. **D.** Same as C, but for the F4 flies. In the F4 generation, larger squares are combined means from both locations, based on n = 41-104.

We infer adaptive tracking when seasonal phenotypic differences in the population persist after four generations of common garden rearing (F4), indicating a genetic basis shaped by seasonal selection rather than environmental effects alone. Indeed, in both the P and F4 generations, time of collection had a significant effect on CT_min_, with the first generation having the lowest CT_min_ (i.e., the most cold tolerant; P generation: LMM, ^2^ = 14.06, df= 2, *P* < 0.001; F4 generation: LMM, ^2^=74.47, df= 2, *P* < 0.001; **Fig. 2C-D)**. In the P generation, CT_min_ was significantly different between the first (4.27 ± 0.1°C) and third time point (5.74 ± 0.1°C; Tukey’s honest significant difference [HSD] *P* < 0.05), while in the F4 generation CT_min_ was significantly different between each of the time points: first (4.98 ± 0.1°C), second (5.51 ± 0.1°C), and third (5.74 ± 0.1°C; Tukey’s HSD, *P* < 0.05). Aggregating across years, CT_min_ increased as a function of Julian date, particularly in the F4 generation (P generation, ANOVA, *F*_1,20_ = 3.88; *P* = 0.062; F4 generation, ANOVA, *F*_1,18_ = 7.34; *P* = 0.014; **Fig. 2B**).

In both the P and F4 generations, when considering weather variables that may be driving evolution of CT_min_, variation in CT_min_ was best predicted by a threshold-based measure of high temperature exposure, specifically the proportion of days exceeding (ρ*_T_*_> 32°C_) during the 0 to 15 days preceding collection, which was negatively associated with CT_min_ at the time of sampling (**Fig. S1**). Notably, while mean temperature showed little association with CT_min_ at short time lags, a weak positive correlation emerged at longer lags (>50 days; **Fig. S2**), potentially reflecting delayed responses to selection across multiple generations, although such patterns could also arise from the strong seasonal autocorrelation of temperature. Overall, this pattern is consistent with seasonal evolution in an environment with coarse-grained variation and the dynamics of adaptive tracking in overwintering species [34], and aligns with model predictions for the rapid evolution of thermal tolerance under fluctuating selection [35].

Together, these results support the hypothesis that adaptive tracking allows *D. suzukii* to dynamically respond to seasonal temperature through evolutionary change in CT_min_. Cold tolerance traits like CT_min_ are quantitative traits shaped by genotype and environment [36], and phenotypic plasticity often plays a larger role in shaping cold tolerance traits than genetic differentiation [37]. Although seasonal shifts in mean CT_min_ may appear modest, they represent rapid evolutionary changes across generations. Indeed, evolutionary rates averaged |*h*| = 0.23 haldanes within growing seasons (**Fig. S3, Table S4**), placing them within the upper range of rates reported for contemporary evolution in natural populations [38]. Thus, even relatively small changes in CT_min_ can reflect substantial evolutionary responses over only a few generations. These rapid seasonal shifts in cold tolerance may be due to fitness trade-offs that are commonly associated with maintenance of cold tolerance [39–41]. *D. suzukii* occurs in a wide range of climates across North America [42, 43] and is present for a large portion of the year at temperate latitudes [44], and therefore adaptive tracking likely provides a mechanism to fine-tune thermal tolerance across seasons. While previous studies have indicated adaptive tracking of other seasonally relevant traits [17, 21, 22, 34, 45], by showing that CT_min_ evolves in a relatively predictable manner over 4 years at two locations, our experiments provide convincing evidence of adaptive tracking of a phenotype relevant for seasonal adaptation. Importantly, these adaptive changes in cold tolerance occurred in a wild population, where gene flow with migration can constrain adaptation [46], further supporting that these changes likely have fitness consequences.

### The complex genetic basis of seasonal adaptive tracking

To identify regions of the genome shaped by seasonal adaptive tracking, we conducted a seasonal contrast analysis (*C*_2_) to identify genomic loci that consistently diverged between the first and last collections across years (**Dataset S2;** see **Figs. S4, S5** for additional seasonality analyses). We assessed significance using a local score approach [47], which identifies genomic regions enriched for *C*_2_ outliers. We identified 65 significant windows (**Fig. 3A, Dataset S3**), collectively spanning 0.1 Mb across the genome, and harboring 3,328 SNPs associated with 57 genes. These windows harbor loci underlying seasonal adaptive tracking, with SNPs enriched in putative regulatory regions, particularly within 5 kb downstream of genes (Odds ratio [OR] = 1.74, *P* = 1.48×10^−18^) and in intergenic regions (OR = 1.66, *P* = 5.78×10^−29^). We visualized genetic variation across these top seasonal loci using PCA, which strongly recapitulated the seasonal sampling scheme and revealed a transition from spring like to fall like genotypes (**Fig. 3B**). PC1 was strongly associated with collection date (**Fig. 3C**; ANOVA, *P* = 0.01), whereas fruit type, year and locality had no significant effects, mirroring the rapid seasonal evolution observed in our trait data.

**Figure 3.**
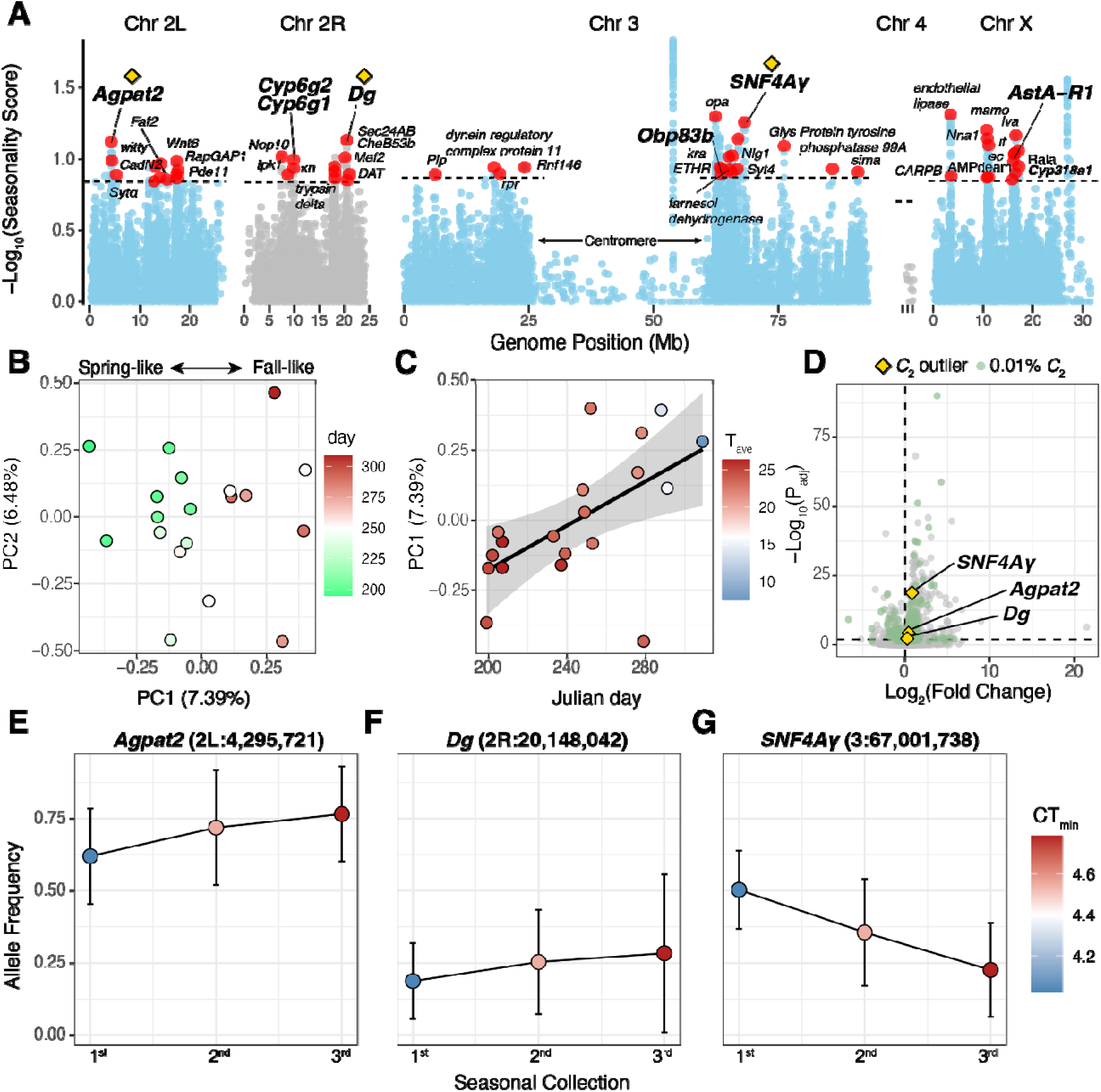
The genetic basis of seasonal adaptive tracking in *D. suzukii*. **A)** Manhattan plot showing the genomic scan for seasonality across the genome of *D. suzukii.* The y-axis shows the Log_10_-transformed Lindley score for the *C*_2_ seasonality statistic calculated using a Lindley parameter ξ = 2 (i.e., “Seasonality score”). The horizontal lines show the significant threshold for enrichment across chromosomes. The red points represent the top *C*_2_ loci within regions of enrichment. **B)** Principal component analysis of outlier loci in the *C*_2_ analysis colored by the date of collection (in Julian date). **C)** PC 1 loading values as a function of the date of collection (in Julian date). Mean temperature during the week of collection is also shown. **D)** Differential expression analysis of an isofemale line from Lexington, KY, in response to a cold-shock exposure. Yellow diamonds indicate genes that harbor at least one outlier mutation from the *C*_2_ outlier analysis. The yellow diamond genes are also indicated in panel A. Green dots represent genes that harbor at least one mutation in the top 0.01% of the *C*_2_ significance distribution. Grey dots harbor no association to the *C*_2_ analysis. **E)** Allele frequencies over collection timepoints for the top seasonal SNP in the *Agpat2* gene, a locus that is differentially expressed in response to cold. The mean phenotypic value for the pooled samples is indicated by color. **F)** Same as E, but for the top SNP in the *Dg* gene. **G)** Same as E, but for the top SNP in the *SNF4A* gene.

Despite the predominance of noncoding SNPs, two missense variants were detected within a tightly clustered set of seasonally fluctuating alleles. Specifically, we observed five top *C*_2_ SNPs within a 140 bp genomic interval spanning *Cyp6g1* and *Cyp6g2*, two genes canonically associated with insecticide resistance [48, 49]. The two nonsynonymous SNPs alter *Cyp6g1* (chr2R:9856476, C→T [V573I]; chr2R:9856487, T→C [K569R]), while two additional variants in the haplotype occur in the 5′ UTR of *Cyp6g2*. Fluctuation in these alleles may reflect seasonal variation in insecticide usage, as maintaining high levels of insecticide resistance can incur trade-offs [50], including possible trade-offs with cold tolerance [18, 51]. Outside of this region, we observed another outlier *C*_2_ variant upstream of the *Odorant-binding protein 83b* (*Obp83b*; chr3:63767198, C→T) gene, a locus involved in the sensory perception of host-associated odors and sensing of sugars [52, 53]. *D. suzukii* can rapidly evolve to new food resources that shift throughout the year [54], and our results indicate there is corresponding evolution of genes involved in finding host fruits.

### Connecting adaptive tracking with the evolution of seasonal cold tolerance

We next characterized connections between genomic adaptive tracking and phenotypic and transcriptional responses to cold stress. We first tested for correlations between allele frequencies and mean cold tolerance among our sequenced Kentucky lines but detected no significant regions of enrichment (**Fig. S4**), reflecting the highly polygenic architecture of cold tolerance. We therefore complemented this analysis by profiling transcriptional responses to a nonlethal cold shock in an isofemale line from Lexington, Kentucky (see *Methods*), identifying 875 differentially expressed genes (*P*_adj_ < 0.01; see ***Supplemental text S3***; **Dataset S4**.) Among these differentially expressed genes, three contained top SNPs from our seasonality analysis (**Fig. 3D**): *1-Acylglycerol-3-phosphate O-acyltransferase 2* (*Agpat2*; *P*_adj_ = 5.78×10^−5^; SNP = chr2L:4,295,721, intronic, T→C; **Fig. 3E**), *Dystroglycan* (*Dg*; *P*_adj_ = 5.00×10^−3^; SNP = chr2R:20,148,042, downstream, G→T; **Fig. 3F**), and *SNF4/AMP-activated protein kinase gamma subunit* (SNF4Aγ; *P*_adj_ = 1.59×10^−19^; SNP = chr3:67,001,738, intronic, C→T; Fig. 3G). All three have established links to metabolic processes important for cold tolerance, including lipid metabolism and membrane remodeling (*Agpat2*), improved mitochondrial respiration at low temperatures (*Dg*), and AMPK mediated metabolic regulation and thermal adaptation (*SNF4A*γ) [55–61]. Together, these loci provide hypothetical links between genomic adaptive tracking and the rapid seasonal evolution of cold tolerance. More generally, our findings show that adaptive tracking targets genes with diverse biological functions (**Table S5**), including those involved in the transcriptional response to cold stress.

### Adaptive tracking is a key contributor to invasiveness

Adaptive tracking, including rapid evolution of cold tolerance, may facilitate *D. suzukii*’s invasion of novel climates [62, 63] and persistence under climate change [11, 64]. As such, we tested whether seasonal SNPs were enriched in genomic regions exhibiting large allele-frequency shifts between native and invasive populations (**Fig. 4A**; [29]). While invasion-associated bottlenecks can drive large frequency shifts at many neutral loci, there is no *a priori* reason to expect these changes to be enriched for adaptive-tracking variants. Seasonal SNPs identified by the *C*_2_ analysis in KY were significantly enriched among SNPs previously associated with invasiveness in North America (OR = 1.26, *P* = 9.48×10^−9^), but significantly under-enriched among those associated with invasiveness in Europe (OR = 0.74, *P* = 1.93×10^−17^). Thus, for North American populations, seasonal adaptive tracking and invasiveness share a genetic basis in North America, but these same alleles are not enriched among the invasiveness alleles in Europe. Interestingly, loci associated with variation in CT_min_ were not enriched among invasion-associated alleles in either continent. This lack of signal is consistent with the polygenic architecture of the trait and suggests that most variants associated with CT_min_ contribute broadly across both native and invaded ranges rather than being necessary for invasion.

**Figure 4:**
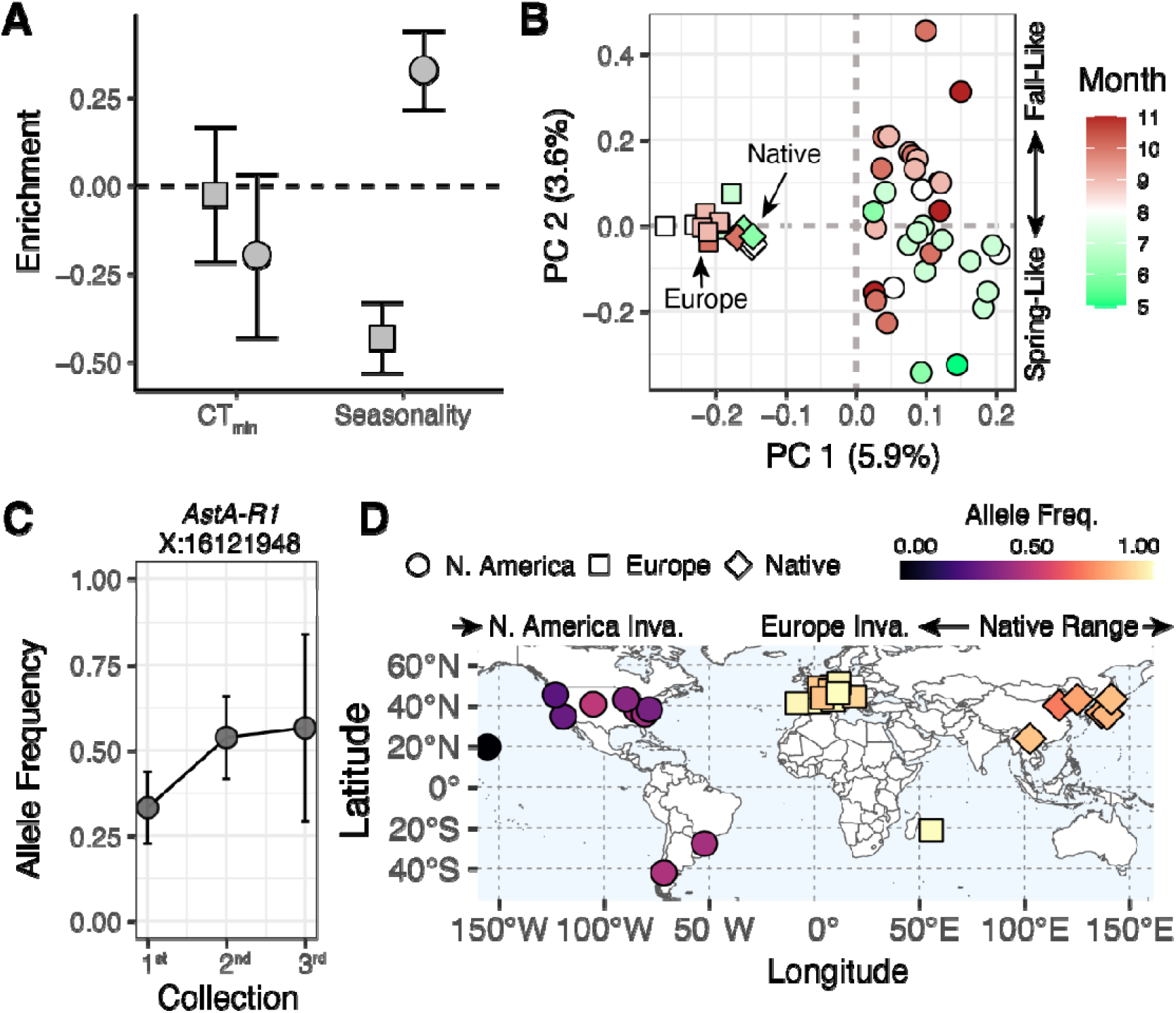
Seasonality and invasiveness share a genetic basis but evolve differently across continents. **A)** Enrichment analyses of the genes associated with invasiveness, both in the Americas and Europe, and two SNP sets characterized in this study (CT_min_ and seasonality [*C*_2_]). Across all panels, circles denote North American data, squares denote European data, and diamonds denote Asian data representing the native range. **(B)** Principal component analysis using the subset of mutations associated with both invasiveness and seasonality. Colors indicate the month of sample collection. Circles represent samples from North America, squares represent samples from Europe, and diamonds represent samples from the native range in Asia. **C)** Allele frequency (mean and 95% confidence intervals) trajectory in KY as a function of field collection time point (i.e., 1^st^, 2^nd^, and 3^rd^, as described in Fig. 2) of a top hit SNP associated with both seasonality, in KY, and invasiveness in North America at the *AstA-R1* locus. **D)** Global allele frequencies for the top hit shown in panel C.

We used a PCA on our global samples to characterize SNPs associated with both adaptive tracking and invasiveness (*n*_SNPs_ = 905; **Fig. 4B**). PC1 (5.9%) captured invasion history (ANOVA, *F*_2,47_ = 226.56, *P* < 2.2×10^−16^)., separating native, European and American populations, whereas PC2 (3.6%) captured seasonal differentiation within the Americas (ANOVA, *F*_1,31_ = 10.31, *P* = 0.0031). Thus, these axes capture invasion history alongside ongoing seasonal adaptation. To illustrate these distinct dynamics, we highlight an intronic variant in the *Allatostatin A receptor 1* gene (*AstA R1*; X-linked:16,121,948, T→C), a locus implicated in sweet sensation in *Drosophila* and mice [65]. This variant exhibits seasonal adaptive tracking in Kentucky (**Fig. 4C**) and marked allele frequency differentiation between invasion routes (**Fig. 4D**), illustrating how seasonal adaptation unfolds against distinct evolutionary histories in Europe and North America.

### Conclusions

Our findings reveal a previously unrecognized link between adaptive tracking and the invasion success of *D. suzukii*. The enrichment of adaptive tracking alleles among variants associated with invasiveness suggests a shared genetic basis, yet the frequencies of these alleles differ between the North American and European invasion fronts. By linking predictable seasonal variation to repeated phenotypic and genomic change across years, our results further support the view that fluctuating selection can drive rapid evolution [17,20]. More importantly, our results identify adaptive tracking as a mechanism that not only promotes persistence in variable environments [38,39], but also as a mechanism that facilitates invasion into novel habitats. More broadly, these findings suggest that adaptive tracking both maintains functional genetic variation and is an important piece of the puzzle underlying invasion success in the Anthropocene [11,27,54]. As environments become increasingly unpredictable, adaptive tracking may play a central role in determining which populations persist, expand, or invade.

## Materials and Methods

### Fly Collection: Locations & Dates

Flies were collected in Fayette and Madison County, Kentucky, beginning in mid-July, when populations of *D. suzukii* started growing [26, 44], and sampling was repeated approximately 6 weeks from the previous collection (4 years collected: 2020-2023; see **Table S1**). Flies were collected by picking infested fruit from the field (blackberries, raspberries, or pokeweed) and allowing flies to emerge from this fruit at room temperature (20°C). Emerging flies were placed on a laboratory diet and moved to a 25°C incubator. Flies that emerged directly from berries were considered the parental (P) generation. After 5 days on lab diet, 20 females were separated to create isofemale lines, while the remaining flies were used for testing thermal tolerance in the parental (P) generation. Isofemale lines were reared at 25°C until the fourth generation (F4). Flies from Virginia, used only in genetic analyses, were collected in Charlottesville, VA. Samples were collected by netting and aspirating twice per year (June and September) across 2017 and 2018.

### Testing Thermal Limits

Flies were tested for thermal tolerance 5-7 days after eclosion. Flies were sorted 2 days before testing by anesthetizing with CO_2_ and given 48 hours at 25 °C to recover. To measure CT_min_, we used a ramping program that started at 25°C for 5 minutes and decreased by −0.25°C per minute until −10°C. Protocols were similar to those described in [66], and are described in full in the supplemental methods. After testing, all flies were stored at −80°C for future genetic analyses.

### DNA Extraction and Sequencing

For the Kentucky samples, we extracted DNA from each time point that had at least 20 P generation flies with equal numbers of males and females (**Table S1**). We pooled flies in groups and extracted DNA with the E.Z.N.A ® Mollusk and DNA Kit #D3373 (Omega Bio-Tek). Library preparation and genomic sequencing were conducted at the Vermont Integrated Genomics Core (VIGR) at the University of Vermont using a Singularity G4 machine (2×250 Paired-End [PE] configuration).

For the Virginia samples, DNA was extracted from pooled samples using a Lithium-Chloride, Potassium-acetate extraction following [13]. We diluted the extracted DNA with water prior to shearing with a Covaris sonicator to create fragments of 500 bp in length. Libraries were prepared using the NEBNext Ultra II kit and dual indices following manufacturer protocols. We performed 8 cycles of PCR during the PCR enrichment step. We quantified the concentration of each library and then pooled the libraries in equal concentrations. After pooling all libraries, we used a Pippen to size-select the DNA for the 600-750 bp range. We then sequenced the pooled library on a NovaSeq with 2×150 PE reads.

### Analyses of Phenotype Data

Analyses were conducted in R version 4.0.5 implemented in RStudio v.1.4.1106. For analysis of CT_min_, we fit full factorial linear mixed models (LMM) using the package ‘lme4’ and analysis of variance (ANOVA) using the package ‘car’. The site of collection was included as a random factor. We first fit a model with time point, sex, year, and generation as fixed variables. We found a strong effect of generation, so we separated the P and F4 generations and fit separate models. All future analyses are done separately for the P and F4 generation Tukey’s honest significant tests (HSD) were performed on these LMMs using the package ‘emmeans’.

We analyzed the association between CT_min_ and environmental variables using regression models that incorporated multiple environmental parameters, including temperature, precipitation, and humidity, each summarized across discretized temporal windows. This approach followed the methods described in Nunez et al. (2024) [12]. Weather data were obtained from the NASA POWER dataset [67], see supplemental for specific variables. We conducted regression analyses using the lme package in R (v.3.1-168). We first generated a null model by regressing CT_min_ against collection site with year and the type of fruit flies were collected from as random effects. We then fit additional models by including one environmental variable as a fixed effect and performed likelihood ratio tests comparing these models with the null model. To assess the overall significance of each model, we conducted one hundred random permutations ([12]; see figure S6 of that paper).

To quantify the lag between the environment and traits or abundance, we conducted a cross-correlation analysis in R with the function ccf from the R package *tseries* [68] with CT_min_ or abundance as the *x*-variable and temperature as the *y*-variable. We also estimated evolutionary rates using the haldane (*h*) metric following [69], based on the change in CT_min_ between consecutive sampling events. Generation number was estimated assuming 16 generations per year during the May-December growing season, and only within-season estimates were included in subsequent analyses.

### Mapping and SNP Calling for Seasonal Samples

DNA reads from all samples (see 53 samples in **Table S2**) were mapped to the *Drosophila suzukii* reference genome assembly (NCBI RefSeq GCF_043229965.1; GenBank GCA_043229965.1), using the latest assembly described in [28]. Prior to mapping, we applied a read level cleaning step to reduce potential contamination from non-target species following the procedure outlined in [70]. Read mapping and post processing were conducted using BWA MEM2 [71] and samtools [72], producing a set of 53 alignment files in bam format.

Variant calling was performed on Pool Seq data from these 53 bam files, comprising of 29 previously published samples representative of worldwide genetic diversity [29, 73] and all newly sequenced samples. SNP discovery was done using the Haplotype Caller implemented in *FreeBayes* [74]. The resulting VCF files were reordered and concatenated using the view and concat programs of *bcftools* [72], and then parsed and transformed into a pooldata object using the *vcf2pooldata()* function of the R package *poolfstat* [31]. The final VCF file was annotated with *SnpEff* [75] using the corresponding NCBI annotation file for the reference assembly. For a targeted subset of SNPs identified in downstream analyses, we also tested whether variants overlapped predicted transcription factor binding motifs using *Tomtom* [76]. See supplemental methods for full cleaning details.

### Spatial and Temporal Demography

To provide a global visualization of the genetic structuring of the populations, we performed a random allele PCA using the *randomallele.pca()* function from the R package *poolfstat* [31]. We further focused on the Kentucky samples by extracting read count data for the 19 corresponding samples, keeping only variants with MAF > 0.01. Pool-Seq based estimators of pairwise *F*_ST_ [77] and hierarchical *F*_ST_ (h*F*_ST_; [78]) were then computed with the *computeFST()* function of *poolfstat* [31]. We tested hypotheses related to the spatial structure and stability of the KY and VA samples using a model-based approach in *moments* [79]. The core methods and principles of these methods, as applied to pool-seq, are described in [80] and [81]. To assess the genomic footprints of overwintering (i.e., boom-and-bust) dynamics, we used the year-to-year *F*_ST_ (calculated using *poolfstat*). See supplemental methods for full description.

### Genomic Scans for Seasonality and Environmental Associations

Genome-wide scan analyses including the *C*_2_ contrast analyses and association with population-specific covariables (i.e.,, CT_min_), were performed with *BayPass* (v3.0; [28, 82]); see supplemental methods. To further identify and delineate significant genomic regions that were enriched for signals of seasonality (based on *C2* seasonal contrast) or CT*_min_* and population-specific covariables, we relied on the local-score approach described in Fariello *et al.* (2017), with ξ = 2 for the underlying Lindley process, and implemented in the R function compute.local.scores() available in the *BayPass* software package (v3.0) [47]. The local score statistic calculates a critical threshold for each chromosome to estimate significance. In cases where no Lindley windows pass the threshold, we considered that no genomic windows were enriched. Invasiveness-associated SNPs were obtained from the published dataset of [29], and genomic regions enriched for invasion-associated alleles were identified using the local score approach.

### Differential Expression Analyses

Molecular responses to cold stress were assessed in a laboratory line of *D. suzukii,* originally collected from infested blackberries in 2017 in Lexington, KY. After collection, a single female was used to start an isofemale line, and this line underwent 20 generations of full-sib inbreeding to establish a line. For the RNA-seq experiment, groups of 10 adult flies, 5-7 days after ecolosion, were exposed to −3°C for 1 h in a refrigerated bath circulator (ThermoFisher Arctic Series) and allowed to recover for 2 h at 25°C prior to flash freezing samples. Control flies were kept at 25°C for duration of the experiment and sampled at the same time as cold-treated flies. RNA was extracted from flies using the Direct-zol RNA Kit (Zymo Research, Irvine, CA), and samples were sent to Novogene for library preparation and sequencing on an Illumina HiSeq 2500. Raw RNA-seq reads were assessed with FastQC [83], trimmed using Trimmomatic [84], and aligned to the NCBI RefSeq GCF_043229965.1 genome with HISAT2 [85]. Differential expression analysis was performed in DESeq2 [86] after filtering lowly expressed genes (<10 counts in ≥3 samples). Gene Ontology enrichment of significant genes (*P*_adj_ < 0.01) was conducted using topGO [87].

## Acknowledgments

We would like to thank the staff at the University of Kentucky Horticultural Research Farm and the Berea College Farm for providing access to field sites, and we also thank Laura Unfried and Leslie Potts for help with sampling. The authors acknowledge the Vermont Advanced Computing Center (VACC; https://www.uvm.edu/vacc) for providing computational resources that contributed to this publication.

## Data Availability

This study incorporates multiple datasets deposited in the National Center for Biotechnology Information (NCBI) repositories, as referenced in the corresponding citations. Data generated for this project is also available in NCBI as PRJNA1420419 (DNA) and PRJNA1526022 (RNA). Supplementary datasets are available in Zenodo (https://zenodo.org/) at: https://doi.org/10.5281/zenodo.18520895. A GitHub repository with all code can be found at: https://github.com/Jcbnunez/ky_swd.git.

## Author Contributions

Conceptualization-E.A.M., N.M.T., J.C.B.N., M.G., K.E.L. Funding acquisition-E.A.M., N.M.T., J.C.B.N., K.E.L Investigation-E.A.M., A.B., M.D., M.G. Formal analysis-E.A.M., J.C.B.N, M.G. K.A.E., M.O.G, A.R.M., S.R., K.E.L Visualization-E.A.M., J.C.B.N, K.A.E., M.O.G, A.R.M., S.R., K.E.L Supervision-N.M.T, J.C.B.N, A.O.B, K.E.L Writing original draft-E.A.M, N.M.T., J.C.B.N., Writing review and editing-E.A.M., M.G., K.A.E., A.B., M.O.G., A.R.M., S.R., A.O.B., K.E.L. J.C.B.N., N.M.T

## Supplemental Methods

### Fly Collection: Locations & Dates

Flies were collected from two sites in central Kentucky, the University of Kentucky Horticultural Farm (37°58D23DN, 84°32D16DW) located in Fayette County, and the Berea College Farm (37°34D06DN, 84°17D12DW) located in Madison County. These farms are 43.5 km apart. Flies were collected early, at the midpoint, and at the end of the growing season to compare thermal tolerance of flies collected at distinct times of year. Specifically, fly collections began in mid-July when populations of D. suzukii started growing in Kentucky [1, 2], and subsequent collections were repeated approximately 6 weeks from the previous collection. The sampling dates for each year can be found in Table S1.

Flies were collected by picking infested fruit from the field. Flies were obtained from either blackberries, raspberries, or pokeweed depending on the date and the site. Fruits were brought back into the lab, put in containers on a layer of vermiculite to soak up excess moisture, and kept on the bench top at room temperature (20°C) until flies emerged. Emerging flies were collected every other day, identified as D. suzukii, placed on a lab diet and moved into a 25°C incubator. The lab diet is the Bloomington Drosophila Stock Center Cornmeal diet, which consists of cornmeal, corn syrup solids, yeast, soy flour, and agar. Flies that emerged directly from berries were considered the parental generation. After 5 days on lab diet, flies were sorted and 20 females were separated to create isofemale lines, while the remaining flies were sorted to be used for the testing thermal tolerance in the parental (P) generation. The 20 females used for the isofemale lines were put in individual vials and allowed to lay eggs for a week, after which the female was discarded. From there each line was reared separately at 25 °C until the fourth generation (F4).

Flies from Virginia were collected at Carter’s Mountain Orchard (37°58D44.51DN, 78°29D23.04DW) located in Charlottesville, VA. Samples were collected twice per year (June and September) across two years (2017 and 2018). Flies were collected using netting and aspiration and identified under a microscope.

### Testing Thermal Limits

All flies used for thermal tests were 5-7 days past eclosion. Flies were separated from either berry enclosures (P generation) or vials (F4 generation) at 1-2 days post eclosion and placed in mixed sex populations on a lab diet in the 25°C incubator for 3 days. On the third day, flies were anesthetized with CO_2_ and sorted by sex. All flies were then put back on lab diet in the 25 °C incubator for 48 hours. For parental generation, 20-25 females and males were separated for testing. For the F4 generation flies 50-70 flies were separated out for testing. However, some flies died or escaped between sorting and final testing, so total numbers for each treatment are in Table S1. There were no P generation flies tested on 8/25/2023 from Berea due to low emergence from fruit.

### Critical Thermal Minimum (CT_min_)

The CT_min_ protocols were similar to those described in Letcheta et al. 2020 [3]. We used programmable refrigerated baths (Thermo Scientific Arctic Series) containing 50% propylene glycol, and these were programmed to cool the liquid at a defined rate. The liquid was pumped through a vertical jacketed column to cool the air in the center [4, 5]. The air temperature in the center of the column was monitored using a type T thermocouple. The ramping program started at 25°C for 5 minutes then decreased by −0.25°C per minute until the temperature reached −10°C. Male and female flies were put into the center of the column, and the column was plugged to keep the flies from escaping. Plastic mesh was inserted into the center of the column to allow flies to perch but still fall through. Once the temperature in the center of the column reached 10°C, the bottom plug was removed. From this point on, any fly that fell out of the column was assumed to have reached its CT_min_. To catch flies that fell from the column, a funnel was attached with a vial of ethanol at the bottom. A new vial was placed underneath after every 0.5°C decrease in temperature. The ramp stopped once all flies fell from the column. Any flies that were obviously stuck to the side of the column or the mesh were not counted. Flies from the vials were counted and sorted by sex, and the CT_min_ was recorded as the maximum temperature from the temperature interval. All flies were then put in one vial and stored at −80°C for future genetic analyses.

### DNA Extraction and Sequencing

For the Kentucky samples, we extracted DNA from parental flies from each time point that had at least 20 flies with equal numbers of males and females. The sampling dates included for DNA extraction and the number of flies in each extraction can be found in Table S1. We also extracted DNA from one F4 generation pool. To extract DNA, we used the E.Z.N.A ® Mollusk and DNA Kit #D3373 from Omega Bio-Tek. We pooled together flies in groups that were equal in sex for extraction. For groups where we sampled more than 30 flies, we split the flies into two separate extractions which were then pooled together in equal DNA concentrations at the end. After extraction, DNA concentration was quantified with a Qubit, and samples were diluted to 10 ug/ul. Library preparation and genomic sequencing were conducted at the Vermont Integrated Genomics Core (VIGR) at the University of Vermont using a Singularity G4 machine (2×250 Paired-End [PE] configuration).

For the Virginia samples, DNA was extracted from pooled samples using a Lithium-Chloride, Potassium-acetate extraction following Bergland et al (2014)[6]. We diluted the extracted DNA with water in a 1:1 ratio prior to shearing with a Covaris sonicator to create fragments of 500 bp in length. Libraries were prepared using the NEBNext Ultra II kit and dual indices following manufacturer protocols. We performed 8 cycles of PCR during the PCR enrichment step. We quantified the concentration of each library and then pooled the libraries in equal concentrations. After pooling all libraries, we used a Pippen to size-select the DNA for the 600-750 bp range. We then sequenced the pooled library on a NovaSeq with 2×150 PE reads.

### Regression Analyses: CT_min_ phenotypic measurements with Julian day

Analyses were conducted in R version 4.0.5 implemented in RStudio Version 1.4.1106. All graphs were made using the package ggplot2. For analysis of CT_min_, we fit full factorial linear mixed models (LMM) using the package ‘lme4’ and analysis of variance (ANOVA) using the package ‘car’. The site of collection was included as a random factor. We first fit a model with all data points with time point, sex, year, and generation as fixed variables. We found a strong effect of generation, so we separated the P and F4 generations and fit separate models. All future analyses are done separately for the P and F4 generation Tukey’s honest significant tests (HSD) were performed on these LMMs using the package ‘emmeans’. We checked that models were normal by diagnostic plots.

### Regression Analyses: Environmental Associations with CT_min_

We analyzed the association between CT_min_ and environmental variables using regression models that incorporated multiple environmental parameters, including temperature, precipitation, and humidity, each summarized across discretized temporal windows. This approach followed the methods described in Nunez et al. (2024) [7]. For every environmental variable, we calculated the minimum, maximum, mean, and variance. For temperature specifically, we also estimated the proportion of days below 5°C (*ρ*_T<5°C_) and the proportion of days above 32°C (*ρ*_T>32°C_). These thresholds represent physiological limits for drosophilids as described by Machado et al. (2021) [8].

Following Nunez et al. (2024), all variables were summarized across 11 windows representing different numbers of days prior to collection: 0 to 7, 0 to 15, 0 to 30, 7 to 15, 15 to 30, 30 to 60, 60 to 90, 15 to 45, 45 to 75, 0 to 60, and 0 to 90. Weather data were obtained from the NASA POWER dataset [9] with a spatial resolution of 0.5° degrees by 0.625° in latitude and longitude. The specific variables used were T2M for temperature (the average air dry bulb temperature at two meters above the surface in °C), PRECTOTCORR for precipitation (a bias corrected estimate of total precipitation at the surface, including the water content in snow, in millimeters per day), and RH2M for relative humidity (the percent ratio of actual to saturation vapor pressure at two meters above the surface). All data were queried at an hourly resolution. Queries used the latitude and longitude associated with the Berea and Lexington sample sites.

We conducted regression analyses using the lme package in R (version). For CT_min_ measurements from both P and F4 generations, we fit two models. The first was a null model in which CT_min_ was regressed on the collection site and year as a random effect, along with the fruit type from which the parental fly was collected, also treated as a random effect. The second model extended the null model by including one environmental variable as a fixed effect. We then performed likelihood ratio tests comparing the two models. To assess the overall significance of each model, we conducted one hundred random permutations following the procedure described in Nunez et al. (2024; see supplementary figure S6 of that paper).

### Cross-Correlation Analysis

To quantify the lag between the environment and traits or abundance, we conducted a cross-correlation analysis in R with the function ccf from the R package tseries [10] with CT_min_ or abundance as the x-variable and temperature as the y-variable.

### Genomic Analyses: Mapping and SNP calling

DNA reads from all samples (see 53 samples in Dataset S2) were mapped to the Drosophila suzukii reference genome assembly (NCBI RefSeq GCF_043229965.1; GenBank GCA_043229965.1), using the latest assembly described in Camus et al. (2025) [11]. Prior to mapping, we applied a read level cleaning step to reduce potential contamination from non-target species following the procedure outlined by Gautier et al. (2023)[12]. Read mapping and post processing were conducted using BWA MEM2 [13] and samtools [14], producing a set of 53 alignment files in bam format (see mapping details in Dataset S2).

Variant calling was performed on Pool Seq data from these 53 bam files, comprising three groups of samples: i) 29 previously published samples representative of worldwide genetic diversity [11, 15, 16] 4 newly sequenced samples collected in Charlottesville, Virginia in 2017 and 2018, and iii) 19 newly sequenced Kentucky samples (P generation) collected in Lexington and Berea, plus one additional pooled sample consisting of an F4 generation of pooled isofemale lines established from Berea (Table S1). SNP discovery was done using the Haplotype Caller implemented in FreeBayes [17] run with the following options: i) −K (to output all alleles that pass input filters, regardless of genotyping outcome, assuming a pooled sequencing model); ii) −C 1 −F 0.01 (requiring at least one count and a fraction of 1% of the counts of the alternate allele to evaluate the position); iii) −G 5 (at least 5 counts supporting an alternate allele in all sample to retain the allele); iv) −-limit-coverage 500 (to downsample per-sample coverage to 500 reads if greater than this coverage); v) −E −1 (to disable clumping of contiguous variants into complex alleles); vi) −n 4 (to evaluate only the best 4 alleles, ranked by sum of supporting quality scores); and vii) −m 30 −q 20 (minimum mapping and base qualities set to 30 and 20, respectively). To improve computational efficiency, analyses were carried out in parallel on a computer grid for 250 kb windows covering the whole genome assembly. The resulting VCF files were reordered and concatenated using the view and concat programs of bcftools [14]. The final VCF file was then parsed and transformed into a pooldata object using the vcf2pooldata() function of the R package poolfstat [18], with the options min.rc = 2, to exclude SNPs with a read count below 2 (indicative of too low coverage), and min.maf = 0.005, to filter out poorly polymorphic sites. An additional filtering step was then applied to remove SNPs with either excessively low or high coverage and insufficient polymorphism, using pooldata.subset with min.cov.per.pool = 5 and cov.qthres.per.pool = c(0, 0.999), and to retain only variants mapping to chrX, chr2L, chr2R, chr3 and chr4 chromosomes. This resulted in a total of 6,299,920 bi-allelic autosomal and 1,167,654 X--linked variants. The final VCF file was annotated with SnpEff [19] using the corresponding NCBI annotation file for the reference assembly. For a targeted subset of SNPs identified in downstream analyses, we also tested whether variants overlapped predicted transcription factor binding motifs using Tomtom [20].

### Genomic Analyses: Spatial and Temporal Demography

To provide a global visualization of the genetic structuring of the populations, we performed a random allele PCA using the randomallele.pca() function of the R package poolfstat[18] on the genomic data. We further focused on the Berea and Lexington samples by extracting read count data for the 19 corresponding samples, keeping only variants with MAF > 0.01. In total, the datasets consisted of 815,578 X-linked and 4,746,028 autosomal SNPs. Pool-Seq based estimators of pairwise F_ST_ [21] and hierarchical F_ST_ (hF_ST_; [22]) were then computed with the computeFST() function of poolfstat [18] run with default options.

We tested hypotheses related to the spatial structure and stability of the KY and VA samples using a model-based approach in moments [23]. The core methods and principles of these methods, as applied to pool-seq, are described in Kapun et al. (2021; DEST 1.0)[24] and Nunez et al. (2025; DEST 2.0)[25]. In brief, we discretized allele frequency estimates using a probabilistic framework weighted by the effective coverage (n_e_; [26–28]) of each pool. These discretized estimates were transformed into folded site frequency spectra (SFS) and used in moments to test three phylogeographic models between all combinations of each KY, and each VA population pools. These models included: quasi-panmixia (i.e., the samples from both KY and VA derived from the same population), symmetrical migration (i.e., the samples from both KY and VA represent two different demes connected by gene-flow; modeled as a single m_i↔ j_ parameter), and asymmetrical migration (i.e., the samples from both KY and VA represent two different demes connected by gene-flow; modeled by two different, m_i→j_ and m_j→i_, parameters). We evaluated model fit across all comparisons using the Akaike Information Criterion (AIC).

To assess temporal demography, particularly the genomic footprints of overwintering (i.e., boom-and-bust) dynamics, we used the year-to-year F_ST_ (calculated using poolfstat). This statistic has been described as a metric of overwintering in D. melanogaster by Nunez et al. (2024) [7] and Nunez et al. (2025) [25]. In brief, the statistic seeks to compare the ratio of genetic differentiation accrued (due to drift) within a growing season (∼10 generations within a year) to that accrued over the few (∼2) generations that correspond to the period of overwintering (i.e., between years). In this context, an increased value for the overwintering F_ST_ is consistent with the action of strong winter bottlenecks. To further explore the patterns of year-to-year F_ST_, we conducted forward genetic simulations using SLiM v.4.3[29]. In brief, we implemented a non-Wright–Fisher simulation modeling the effects of selection from seasonal fluctuations in the form of cyclic population crashes. We modeled a single population of 5,500 individuals with a 100 kb genome containing neutral mutations imported from a neutral VCF generated from msprime [30]. A single neutral mutation type was applied uniformly across the chromosome, with a recombination rate of 1.25×10⁻⁷ per base per generation and a mutation rate of 1×10⁻⁹. Reproduction followed a Poisson distribution with mean R = 2.3, and generations were non-overlapping, with adults removed at the start of each cycle. Fitness Scaling was determined by density dependence and fitness scaling modified by an environment-dependent performance curve following a modified Norberg function Norberg (2004) [31]:

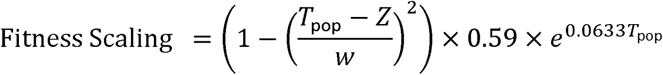

In this function, T_pop_ is the subpopulation’s current environment, “Z” is the optimum environment (0.0), and “w” controls the width of the performance curve. Environment is set at the start of each cycle alternating between two deterministic states: 12 generations of optimal environment (T_pop_ = 0.0) and 4-generations of non-optimal conditions (T_pop_ = 0.1 to .9) with larger values of T_pop_ driving greater population crashes. At each generation, the simulation output population size, time (generations past), and allele frequencies for all alleles in the genome for 10 replicates of each value of T_pop_.

All simulation outputs from SLiM were processed and analyzed in R. Allele frequency data were imported from multiple output files and concatenated into a single dataset. Low-frequency variants (minor allele frequency <0.05) and near-fixation variants (>0.95) were filtered out. Population size trajectories were visualized by summarizing the mean number of individuals per generation across replicates and conditions. Harmonic mean population sizes (N_e_) were calculated to capture the effects of demographic fluctuations within and across seasonal periods. To assess temporal shifts in allele frequencies, pairwise F_ST_ was calculated using the ANOVA method implemented in poolfstat. Artificial coverage was held constant to standardize inputs, and pool sizes were taken from mean population sizes at each generation. Comparisons were summarized within and across both seasons and years.

### Genomic Analyses: Genomic Scans for Seasonality and Environmental Associations

Genome-wide scan analyses, including X^T^X scan for adaptive differentiation, C_2_ contrast analyses and association with population-specific covariable (CT_min_ and ρ*_T_*_> 32°C_ were performed with BayPass (v3.0; [11, 32]). We first extracted read count data for the 19 Berea and Lexington samples from the complete pooldata object on the 53 samples (see above), keeping only variants with MAF>0.01 and transformed the obtained object into BayPass input files using the pooldata2genobaypass()function in poolfstat. In total, the datasets consisted of 815,578 X-linked and 4,746,028 autosomal SNPs. The autosomal and X chromosome data were analyzed separately default BayPass option to estimate the scaled covariance matrix of population allelic frequencies (Ω), accounting for the neutral population structure, the overall differentiation (with the X^T^X statistic), the seasonal contrast (with the C statistic), and the support for association with CT_min_ and ρ*_T_*_> 32°C_ (with BF estimated with the default Importance Sampling algorithm). To further identify and delineate significant genomic regions that are enriched for signals of seasonality (based on C2 seasonal contrast) or CTmin and population-specific covariables, we relied on the local-score approach described in Fariello et al. (2017) and implemented in the R function compute.local.scores available in the BayPass software package (v3.0), using default options (see details in BayPass manual). The local score statistic calculates a critical threshold for each chromosome to estimate significance. In cases where no Lindley windows pass the threshold, we considered that no genomic windows were enriched.

### Transcriptomics Analyses: Differential Expression

Quality control on raw reads was performed on the raw reads using FastQC v0.12.1 [33] and a MultiQC v1.15 [34] report of the FastQC output was generated with all samples having Phred quality scores above 30. Reads were preprocessed and filtered using Trimmomatic v0.39 [35] for paired-end reads. Minimum base quality for starts and ends of reads were set to a minimum of 3, roughly equal to an error rate of 0.5. After initial trimming, any 4-nucleotide window below 95% certainty was trimmed. Any read shorter than 36 nucleotides were trimmed. Quality control was performed on the trimmed reads with FastQC and a MultiQC report was generated. MultiQC revealed trimming had little impact, indicating little degradation or adapter contamination on raw dataset. We used the hisat2 v2.2.1 [36] package to map reads to the reference genome. BAM files were sorted using Samtools v1.19.2 [37]. Qualimap v2.3 [38] was used to ensure the quality of all files. Using the featurecounts function from subread v2.0.8 [39], we created a gene count matrix based on GTF annotation from NCBI RefSeq GCF_043229965.1. Using the DESeq2 v1.50.2 [40] package, we then generated a DESeq2 object from the gene count matrix using the control group as baseline level. The dataset was filtered to remove any genes with fewer than 10 reads in at least 3 samples before running differential expression analysis. The expression table was filtered for genes with an adjusted p-value under 0.01. Using NCBI GO annotation, differentially expressed genes without GO annotations were removed from analysis. Using the topGO package v2.62.0 [41], GO enrichment was performed using a parent-child algorithm and Fisher’s exact test. The results were converted to a table and filtered for annotations associated with between 5 and 500 genes. The top 20 enriched GO terms were plotted. We then compared this data with GO enrichment performed on genes known to be involved with seasonality.

---

**Table S1.**
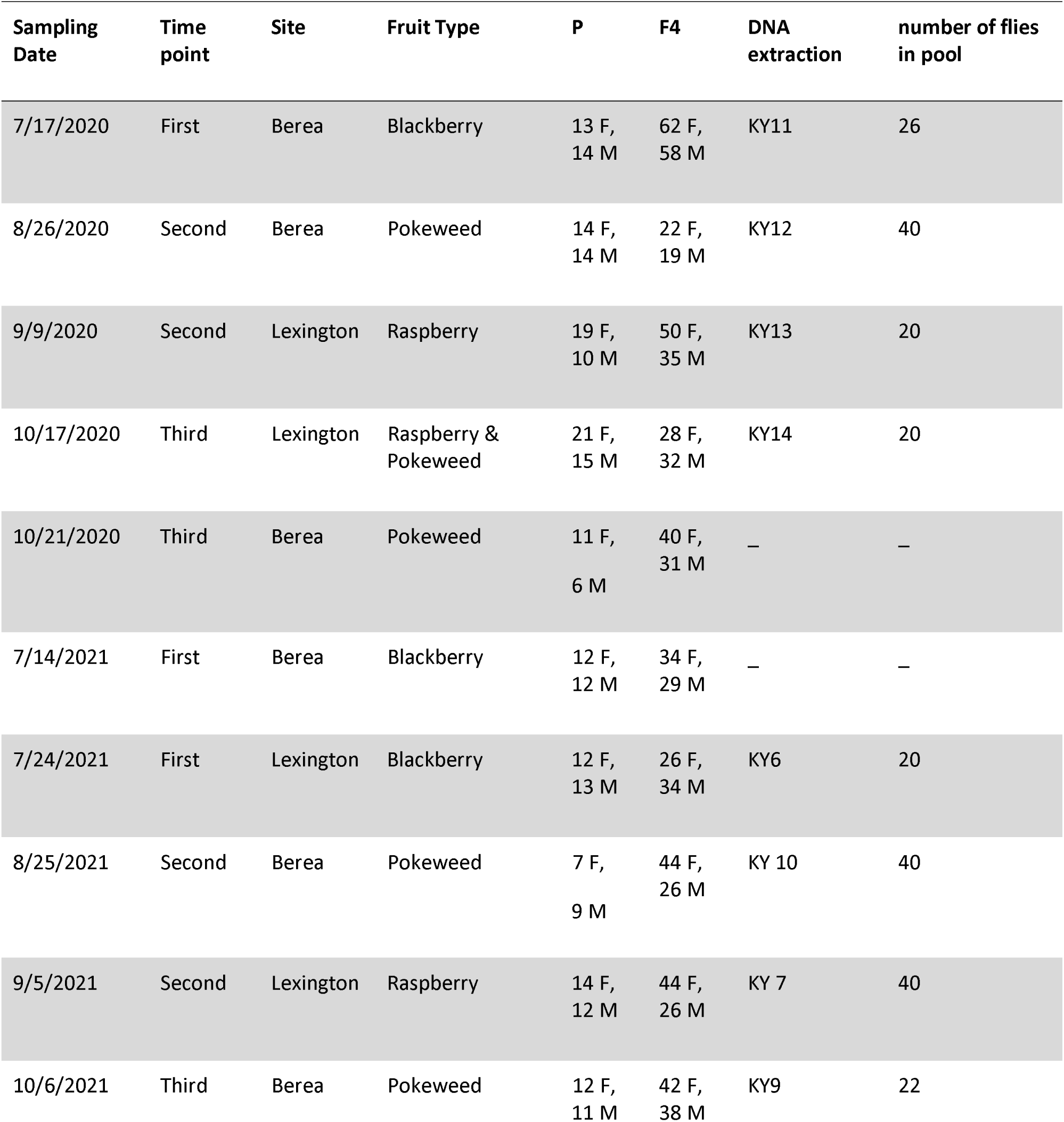
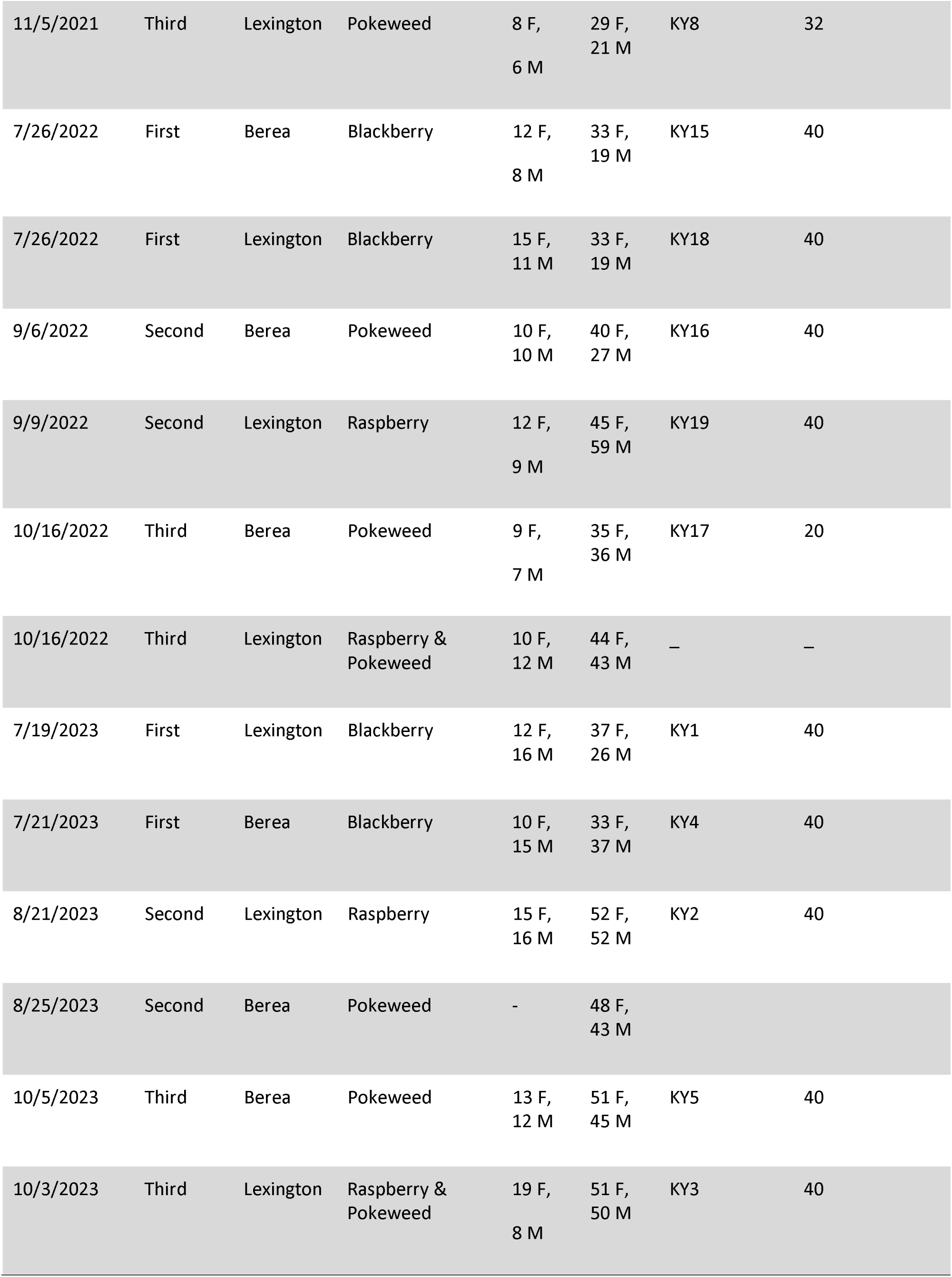
Full information on sampling and testing for Kentucky flies. Columns indicate sampling date, time point, location, fruit the flies were originally reared from, the number of flies tested for CT_min_ in the P and F4 generation, which samples were used in DNA extraction, and the number of flies in the DNA pool.

| Sampling Date | Time point | Site | Fruit Type | P | F4 | DNA extraction | number of flies in pool |
|---|---|---|---|---|---|---|---|
| 7/17/2020 | First | Berea | Blackberry | 13 F, 14 M | 62 F, 58 M | KY11 | 26 |
| 8/26/2020 | Second | Berea | Pokeweed | 14 F, 14 M | 22 F, 19 M | KY12 | 40 |
| 9/9/2020 | Second | Lexington | Raspberry | 19 F, 10 M | 50 F, 35 M | KY13 | 20 |
| 10/17/2020 | Third | Lexington | Raspberry & Pokeweed | 21 F, 15 M | 28 F, 32 M | KY14 | 20 |
| 10/21/2020 | Third | Berea | Pokeweed | 11 F,<br>6 M | 40 F, 31 M | _ | _ |
| 7/14/2021 | First | Berea | Blackberry | 12 F, 12 M | 34 F, 29 M | _ | _ |
| 7/24/2021 | First | Lexington | Blackberry | 12 F, 13 M | 26 F, 34 M | KY6 | 20 |
| 8/25/2021 | Second | Berea | Pokeweed | 7 F,<br>9 M | 44 F, 26 M | KY 10 | 40 |
| 9/5/2021 | Second | Lexington | Raspberry | 14 F, 12 M | 44 F, 26 M | KY 7 | 40 |
| 10/6/2021 | Third | Berea | Pokeweed | 12 F, 11 M | 42 F, 38 M | KY9 | 22 |
| 11/5/2021 | Third | Lexington | Pokeweed | 8 F,<br>6 M | 29 F, 21 M | KY8 | 32 |
| 7/26/2022 | First | Berea | Blackberry | 12 F,<br>8 M | 33 F, 19 M | KY15 | 40 |
| 7/26/2022 | First | Lexington | Blackberry | 15 F, 11 M | 33 F, 19 M | KY18 | 40 |
| 9/6/2022 | Second | Berea | Pokeweed | 10 F, 10 M | 40 F, 27 M | KY16 | 40 |
| 9/9/2022 | Second | Lexington | Raspberry | 12 F,<br>9 M | 45 F, 59 M | KY19 | 40 |
| 10/16/2022 | Third | Berea | Pokeweed | 9 F,<br>7 M | 35 F, 36 M | KY17 | 20 |
| 10/16/2022 | Third | Lexington | Raspberry & Pokeweed | 10 F, 12 M | 44 F, 43 M | _ | _ |
| 7/19/2023 | First | Lexington | Blackberry | 12 F, 16 M | 37 F, 26 M | KY1 | 40 |
| 7/21/2023 | First | Berea | Blackberry | 10 F, 15 M | 33 F, 37 M | KY4 | 40 |
| 8/21/2023 | Second | Lexington | Raspberry | 15 F, 16 M | 52 F, 52 M | KY2 | 40 |
| 8/25/2023 | Second | Berea | Pokeweed | - | 48 F, 43 M |  |  |
| 10/5/2023 | Third | Berea | Pokeweed | 13 F, 12 M | 51 F, 45 M | KY5 | 40 |
| 10/3/2023 | Third | Lexington | Raspberry & Pokeweed | 19 F,<br>8 M | 51 F, 50 M | KY3 | 40 |

**Table S2:** Metadata for all samples used in this study. Sample Id orig: indicated the name of the samples used in the BAM file creation; Lat: Latitude; long: Longitude; continent: continent of origin; country: country of origin; city: city of origin; Collection date: date when samples were collected from the field. Fruit type: if flies were obtained directly from fruit, which cultivar was sampled; nFlies: Number of flies included in the pool; nmales: Number of males included in the pool. Fly type: Type of line (wild-caught, isofemale, F4); Sampling strategy: whther samples were caught using traps or directly sampled from fruit as emergent larvae; Seq platform: Sequencing platform; Range: Native or invasive range; SRA: NCBI SRA accession; Reference: Published reference.

| Sample Id orig | lat | long | continent | country | city | Collection date | Fruit type | nFlies | nmales | Fly type | Sampling strategy | Seq platform | Range | SRA Accession | reference |
|---|---|---|---|---|---|---|---|---|---|---|---|---|---|---|---|
| **BR-Pal** | -27.72 | -52.17 | South America | Brazil | Porto Alegre | 2014-07 | NA | 50 | 33 | wild | trap | HS2500(PE125) | Invasive AME | SRR10260033 | Olazcuaga et al. (2020) |
| **CN-Bei** | 40 | 116.35 | Asia | China | Beijing | 2014-06 | NA | 50 | 11 | wild | trap | HS2500(PE125) | NAT | SRR10260017 | Olazcuaga et al. (2020) |
| **CN-Lia** | 42.96 | 125.09 | Asia | China | Liaoyuan city | 2014-08 | NA | 50 | 40 | wild | trap | HS2500(PE125) | NAT | SRR10260016 | Olazcuaga et al. (2020) |
| **CN-Shi** | 23.7 | 102.48 | Asia | China | Shiping | 2014-06;08 | NA | 50 | 47 | wild | trap | HS2500(PE125) | NAT | SRR10260024 | Olazcuaga et al. (2020) |
| **DE-Dos** | 49.45 | 8.66 | Europe | Germany | Dossenheim | 2015-08 | NA | 50 | 42 | wild | trap | HS2500(PE125) | Invasive EUR | SRR10260021 | Olazcuaga et al. (2020) |
| **ES-Bar** | 41.36 | 1.96 | Europe | Spain | Barcelona | 2014-07 | NA | 50 | 50 | wild | trap | HS2500(PE125) | Invasive EUR | SRR10260019 | Olazcuaga et al. (2020) |
| **FR-Cor** | 42.35 | 9 | Europe | France | Lozzi | 2016-08 | NA | 50 | 25 | wild | trap | HS2500(PE125) | Invasive EUR | SRR10260018 | Olazcuaga et al. (2020) |
| **FR-Lez** | 43.7 | 3.83 | Europe | France | Montpellier | 2014-07 | NA | 100 | 50 | wild | trap | HS2500(PE125) | Invasive EUR | SRR10260015 | Olazcuaga et al. (2020) |
| **FR-Par** | 48.84 | 2.36 | Europe | France | Paris | 2016-11 | NA | 100 | 50 | wild | trap | HS2500(PE125) | Invasive EUR | SRR10260014 | Olazcuaga et al. (2020) |
| **FR-Run** | -21.15 | 55.64 | Africa | France | La Reunion Island | 2016-09 | NA | 100 | 50 | wild | trap | HS2500(PE125) | Invasive EUR | SRR10260013 | Olazcuaga et al. (2020) |
| **IT-Tre** | 46.04 | 11.15 | Europe | Italy | Trento | 2014-09 | NA | 100 | 60 | wild | trap | HS2500(PE125) | Invasive EUR | SRR10260034 | Olazcuaga et al. (2020) |
| **JP-Kan** | 36.5 | 136.6 | Asia | Japan | Kanazawa | 2019-10 | NA | 212 | 183 | wild | trap | NS6000(PE150) | NAT | SRR24598921 | Feng et al. (2024) |
| **JP-Sap2** | 43.11 | 141.24 | Asia | Japan | Sapporo | 2019-08-XX | NA | 151 | 127 | wild | trap | NS6000(PE150) | NAT | SRR24598920 | Feng et al. (2024) |
| **JP-Sap** | 43.05 | 141.35 | Asia | Japan | Sapporo | 2014-07;2016-09 | NA | 50 | 46 | wild | trap | HS2500(PE125) | NAT | SRR10260023 | Olazcuaga et al. (2020) |
| **JP-Tok** | 35.64 | 139.37 | Asia | Japan | Tokyo | 2016-06 | NA | 50 | 10 | wild | trap | HS2500(PE125) | NAT | SRR10260022 | Olazcuaga et al. (2020) |
| **KY10** | 37.5683 | -84.2867 | North America | USA | Berea | 8/25/21 | Pokeweed | 40 | 20 | wild | Fruit sample | G4 | Invasive AME | SRR37160421 | This Study |
| **KY11** | 37.5683 | -84.2867 | North America | USA | Berea | 7/17/20 | Blackberry | 26 | 13 | wild | Fruit sample | G4 | Invasive AME | SRR37160410 | This Study |
| **KY12** | 37.5683 | -84.2867 | North America | USA | Berea | 8/26/20 | Pokeweed | 40 | 20 | wild | Fruit sample | G4 | Invasive AME | SRR37160409 | This Study |
| **KY13** | 37.9731 | -84.5378 | North America | USA | Lexington | 9/9/20 | Raspberry | 20 | 10 | wild | Fruit sample | G4 | Invasive AME | SRR37160408 | This Study |
| **KY14** | 37.9731 | -84.5378 | North America | USA | Lexington | 10/17/20 | Pokeweed/Raspberry | 20 | 10 | wild | Fruit sample | G4 | Invasive AME | SRR37160407 | This Study |
| **KY15** | 37.5683 | -84.2867 | North America | USA | Berea | 7/26/22 | Blackberry | 40 | 20 | wild | Fruit sample | G4 | Invasive AME | SRR37160406 | This Study |
| **KY16** | 37.5683 | -84.2867 | North America | USA | Berea | 9/6/22 | Pokeweed | 40 | 20 | wild | Fruit sample | G4 | Invasive AME | SRR37160405 | This Study |
| **KY17** | 37.5683 | -84.2867 | North America | USA | Berea | 10/16/22 | Pokeweed | 20 | 10 | wild | Fruit sample | G4 | Invasive AME | SRR37160404 | This Study |
| **KY18** | 37.9731 | -84.5378 | North America | USA | Lexington | 7/26/22 | Blackberry | 40 | 20 | wild | Fruit sample | G4 | Invasive AME | SRR37160403 | This Study |
| **KY19** | 37.9731 | -84.5378 | North America | USA | Lexington | 9/9/22 | Raspberry | 40 | 20 | wild | Fruit sample | G4 | Invasive AME | SRR37160420 | This Study |
| **KY1** | 37.5683 | -84.2867 | North America | USA | Lexington | 7/19/23 | Blackberry | 40 | 20 | wild | Fruit sample | G4 | Invasive AME | SRR37160422 | This Study |
| **KY20** | 37.5683 | -84.2867 | North America | USA | Berea | 7/21/23 | Blackberry | 40 | 20 | F4 | Fruit sample | G4 | Invasive AME | SRR37160418 | This Study |
| **KY2** | 37.9731 | -84.5378 | North America | USA | Lexington | 8/21/23 | Raspberry | 40 | 20 | wild | Fruit sample | G4 | Invasive AME | SRR37160419 | This Study |
| **KY3** | 37.9731 | -84.5378 | North America | USA | Lexington | 10/3/23 | Pokeweed/Raspberry | 40 | 20 | wild | Fruit sample | G4 | Invasive AME | SRR37160417 | This Study |
| **KY4** | 37.5683 | -84.2867 | North America | USA | Berea | 7/21/23 | Blackberry | 40 | 20 | wild | Fruit sample | G4 | Invasive AME | SRR37160416 | This Study |
| **KY5** | 37.5683 | -84.2867 | North America | USA | Berea | 10/5/23 | Pokeweed | 40 | 20 | wild | Fruit sample | G4 | Invasive AME | SRR37160415 | This Study |
| **KY6** | 37.9731 | -84.5378 | North America | USA | Lexington | 7/24/21 | Blackberry | 20 | 10 | wild | Fruit sample | G4 | Invasive AME | SRR37160414 | This Study |
| **KY7** | 37.9731 | -84.5378 | North America | USA | Lexington | 9/5/21 | Raspberry | 40 | 20 | wild | Fruit sample | G4 | Invasive AME | SRR37160413 | This Study |
| **KY8** | 37.9731 | -84.5378 | North America | USA | Lexington | 11/5/21 | Pokeweed | 32 | 16 | wild | Fruit sample | G4 | Invasive AME | SRR37160412 | This Study |
| **KY9** | 37.5683 | -84.2867 | North America | USA | Berea | 7/14/21 | Blackberry | 22 | 11 | wild | Fruit sample | G4 | Invasive AME | SRR37160411 | This Study |
| **AR-Gol** | -42.02 | -71.52 | South America | Argentina | Golondrinas | 2023-02 | NA | 100 | 100 | wild | trap | NS6000(PE150) | Invasive AME | SRR29639537 | Camus et al. (2024) |
| **CH-Del** | 47.37 | 7.35 | Europe | Switzerland | Delemont | 2014-10 | NA | 58 | 30 | wild | trap | NS6000(PE250) | Invasive EUR | SRR26544765 | Camus et al. (2024) |
| **DE-Jen** | 50.93 | 11.56 | Europe | Germany | Jena | 2016-09 | NA | 100 | 0 | wild | trap | NS6000(PE250) | Invasive EUR | SRR29639538 | Camus et al. (2024) |
| **RS-Zaj** | 44.45 | 19.25 | Europe | Serbia | Zaja ca | 2020-09 | NA | 100 | 100 | wild | trap | NS6000(PE250) | Invasive EUR | SRR26544764 | Camus et al. (2024) |
| **PT-CP21** | 41.054 | -8.28 | Europe | Portugal | Castelo de Paiva | 2021-09-XX | NA | 80 | 40 | wild | trap | NA | Invasive EUR | SRR28145941 | Sario et al. (2024) |
| **US-CAOx** | 34.15 | -119.13 | North America | USA | Oxnard | 2018-11 | NA | 100 | 0 | wild | trap | NS6000(PE150) | Invasive AME | SRR24598919 | Feng et al. (2024) |
| **US-Col** | 40.57 | -105.09 | North America | USA | Fort Collins | 2015-09 | NA | 50 | 26 | wild | trap | HS2500(PE125) | Invasive AME | SRR10260032 | Olazcuaga et al. (2020) |
| **US-Haw** | 19.67 | -155.47 | North America | USA | Hilo | 2016-06 | NA | 50 | 25 | wild | trap | HS2500(PE125) | Invasive AME | SRR10260031 | Olazcuaga et al. (2020) |
| **US-Nca** | 35.7 | -80.62 | North America | USA | Raleigh | 2016-10 | NA | 100 | 50 | wild | trap | HS2500(PE125) | Invasive AME | SRR10260030 | Olazcuaga et al. (2020) |
| **US-Sdi** | 32.72 | -117.16 | North America | USA | San Diego | 2014-05 | NA | 50 | 32 | wild | trap | HS2500(PE125) | Invasive AME | SRR10260029 | Olazcuaga et al. (2020) |
| **US-Sok** | 45.22 | -123.07 | North America | USA | Dayton | 2014-10 | NA | 75 | 55 | wild | trap | HS2500(PE125) | Invasive AME | SRR10260028 | Olazcuaga et al. (2020) |
| **US-Vir17-VA1** | 37.99 | -78.47 | North America | USA | Charlottesville | 7/20/17 | FieldCollected (Peach/Apple) | 36 | 11 | wild | trap | NS6000(PE150) | Invasive AME | SRR14476475 | This Study |
| **US-Vir17-VA2** | 37.99 | -78.47 | North America | USA | Charlottesville | 9/28/17 | FieldCollected (Peach/Apple) | 37 | 24 | wild | trap | NS6000(PE150) | Invasive AME | SRR14476474 | This Study |
| **US-Vir18-VA3** | 37.99 | -78.47 | North America | USA | Charlottesville | 7/19/18 | FieldCollected (Peach/Apple) | 32 | 16 | wild | trap | NS6000(PE150) | Invasive AME | SRR14476473 | This Study |
| **US-Vir18-VA4** | 37.99 | -78.47 | North America | USA | Charlottesville | 9/6/18 | FieldCollected (Peach/Apple) | 46 | 40 | wild | trap | NS6000(PE150) | Invasive AME | SRR14476472 | This Study |
| **US-Wat** | 36.9 | -121.75 | North America | USA | Watsonville | 2014-10 | NA | 50 | 46 | wild | trap | HS2500(PE125) | Invasive AME | SRR10260026 | Olazcuaga et al. (2020) |
| **US-Wis2** | 42.91 | -89.44 | North America | USA | Oregon | 2018-06-XX::2018-08-XX | NA | 109 | 0 | wild | trap | NS6000(PE150) | Invasive AME | SRR24598918 | Feng et al. (2024) |
| **US-Wis** | 42.97 | -89.69 | North America | USA | Barneveld | 2016-11 | NA | 75 | 30 | wild | trap | HS2500(PE125) | Invasive AME | SRR10260025 | Olazcuaga et al. (2020) |

**Table S3:** Realized coverage of all the Whole-Genome Sequence samples. Mean coverages were estimated with mosdepth [v0.3.6; Pedersen and Quinlan (2017)] using the bam files obtained from the mapping of the filtered raw reads on the new IsoJap1 chromosome-level assembly (Camus et al. 2024) complemented with the complete mitochondrial sequence of D. suzukii (GenBank Accession ID: KU588141) and the complete sequence for the Wolbachia endosymbiont (wRI strain; NCBI ID: GCA 000022285); for the i) autosomes (A) consisting of chr2L, chr2R, chr3, and chr4 chromosomes; ii) X-chromosome; iii) Y chromosome; iv) the Mitochondria (M), and v) Wolbachia (W). Note that coverages for Mito and wRI is null for all KY samples since mapping was carried out on the IsoJap1 without the Mito and wRI sequences.

| Id | A | X | Y | M | W |
|---|---|---|---|---|---|
| **AR-Gol** | 53.17 | 27.05 | 14.96 | 13229.8 | 0.02 |
| **BR-Pal** | 62.78 | 47.37 | 6.54 | 1598.92 | 9.23 |
| **CH-Del** | 61.58 | 49.23 | 8.54 | 12101.2 | 5.33 |
| **CN-Bei** | 82.65 | 76.69 | 4.23 | 2580.44 | 61.93 |
| **CN-Lia** | 59.2 | 37.67 | 9.24 | 7270.4 | 40.02 |
| **CN-Shi** | 58.56 | 31.76 | 12.08 | 5905.37 | 48.25 |
| **DE-Dos** | 70.37 | 44.9 | 13.44 | 46545.2 | 35.37 |
| **DE-Jen** | 50.26 | 49.6 | 0.92 | 11254.4 | 8.73 |
| **ES-Bar** | 66.56 | 34.11 | 17.24 | 6193.04 | 11.09 |
| **FR-Cor** | 60.89 | 51.92 | 4.77 | 3454.83 | 16.76 |
| **FR-Lez** | 74.03 | 58.43 | 6.95 | 2275.75 | 11.21 |
| **FR-Par** | 61.33 | 49.24 | 7.45 | 4624.78 | 2.31 |
| **FR-Run** | 78.15 | 60.83 | 7.02 | 5327.58 | 23.14 |
| **IT-Tre** | 60.34 | 44.51 | 9.13 | 21602 | 12.83 |
| **JP-Kan** | 105.55 | 71.61 | 19.66 | 25108.4 | 21.93 |
| **JP-Sap** | 73.23 | 37.49 | 16.78 | 9082.81 | 4.26 |
| **JP-Sap2** | 96.67 | 62.41 | 16.67 | 6072.55 | 6.39 |
| **JP-Tok** | 57.75 | 51.23 | 3.18 | 7773.94 | 14.99 |
| **KY1** | 39.84 | 33.7 | 3.16 | NA | NA |
| **KY10** | 33.88 | 28.23 | 2.64 | NA | NA |
| **KY11** | 29.18 | 22.2 | 2.58 | NA | NA |
| **KY12** | 27.63 | 20.84 | 2.83 | NA | NA |
| **KY13** | 28.19 | 21.9 | 2.75 | NA | NA |
| **KY14** | 32.67 | 25.77 | 2.73 | NA | NA |
| **KY15** | 34.86 | 31.74 | 2.07 | NA | NA |
| **KY16** | 30.04 | 25.48 | 2.51 | NA | NA |
| **KY17** | 32.41 | 27.57 | 2.71 | NA | NA |
| **KY18** | 28.94 | 23.96 | 2.21 | NA | NA |
| **KY19** | 35.7 | 30.15 | 3.05 | NA | NA |
| **KY2** | 36.97 | 30.34 | 3.37 | NA | NA |
| **KY20** | 38.57 | 32.82 | 2.83 | NA | NA |
| **KY3** | 33.97 | 29.44 | 3.73 | NA | NA |
| **KY4** | 35.6 | 29.18 | 2.84 | NA | NA |
| **KY5** | 35.58 | 30.56 | 2.59 | NA | NA |
| **KY6** | 30.85 | 27.01 | 1.99 | NA | NA |
| **KY7** | 39.31 | 32.03 | 3.18 | NA | NA |
| **KY8** | 37.98 | 31.56 | 3.1 | NA | NA |
| **KY9** | 33.12 | 29.37 | 2.43 | NA | NA |
| **PT-CP21** | 81.9 | 65.06 | 9.14 | 13362.2 | 2.58 |
| **RS-Zaj** | 45.8 | 23.31 | 13.32 | 6757.43 | 2.11 |
| **US-CAOx** | 42.94 | 43.71 | 0.43 | 9780.01 | 0.34 |
| **US-Col** | 66.83 | 52.55 | 6.55 | 14251.1 | 1.12 |
| **US-Haw** | 79.46 | 62.12 | 6.27 | 7756.25 | 1.53 |
| **US-Nca** | 61.05 | 48.01 | 5.38 | 4326.44 | 3.65 |
| **US-Sdi** | 73.79 | 54.28 | 7.61 | 7085.77 | 2.8 |
| **US-Sok** | 54.75 | 34.18 | 10.76 | 11429.8 | 12.83 |
| **US-Vir17-VA1** | 315.19 | 271.06 | 29.58 | 5269.5 | 2.76 |
| **US-Vir17-VA2** | 26.34 | 17.85 | 4.44 | 1189.08 | 0.95 |
| **US-Vir18-VA3** | 23.9 | 18.23 | 3.04 | 1750.32 | 1.21 |
| **US-Vir18-VA4** | 94.51 | 54.38 | 24.41 | 389.83 | 4.85 |
| **US-Wat** | 59.69 | 33.56 | 10.64 | 5104.49 | 15.39 |
| **US-Wis** | 65.67 | 52.06 | 6.33 | 12957.8 | 0.41 |
| **US-Wis2** | 33.6 | 32.85 | 0.4 | 18360.9 | 1.08 |

**Table S4:** Phenotypic values (CT_min_) for D. suzukii collections used in the evolutionary rate (i.e., haldane [h]) analysis.

| *Locality* | *Mean CT<sub>min</sub>* | *SD CT<sub>min</sub>* | *Sequenced pool* | *Date* | *N* |
|---|---|---|---|---|---|
| **Berea** | 4.68 | 2.5724097 | KY4 | 7/21/23 | 25 |
| **Berea** | 4.17 | 1.3360389 | KY5 | 10/5/23 | 25 |
| **Berea** | 5.44167 | 1.5830396 | KY9 | 7/14/21 | 23 |
| **Berea** | 4.23125 | 1.4884976 | KY10 | 8/25/21 | 16 |
| **Berea** | 4.24444 | 2.6243436 | KY11 | 7/17/20 | 27 |
| **Berea** | 3.97931 | 2.6909079 | KY12 | 8/26/20 | 28 |
| **Berea** | 3.77407 | 0.9062147 | KY15 | 7/26/22 | 20 |
| **Berea** | 4.825 | 1.1795963 | KY16 | 9/6/22 | 20 |
| **Berea** | 5.23438 | 1.3767981 | KY17 | 10/16/22 | 16 |
| **Lexington** | 3.82143 | 1.1443443 | KY1 | 7/19/23 | 28 |
| **Lexington** | 4.95968 | 1.6420221 | KY2 | 8/21/23 | 31 |
| **Lexington** | 6.02778 | 1.3180211 | KY3 | 10/3/23 | 27 |
| **Lexington** | 4.54 | 1.7061164 | KY6 | 7/24/21 | 25 |
| **Lexington** | 3.66154 | 1.363254 | KY7 | 9/5/21 | 26 |
| **Lexington** | 4.77143 | 1.4917355 | KY8 | 11/5/21 | 14 |
| **Lexington** | 6.51035 | 1.7899212 | KY13 | 9/9/20 | 29 |
| **Lexington** | 4.61944 | 1.9586904 | KY14 | 10/17/20 | 36 |
| **Lexington** | 3.08636 | 0.7389152 | KY18 | 7/26/22 | 26 |
| **Lexington** | 3.67619 | 1.2497619 | KY19 | 9/9/22 | 21 |

**Table S5:**
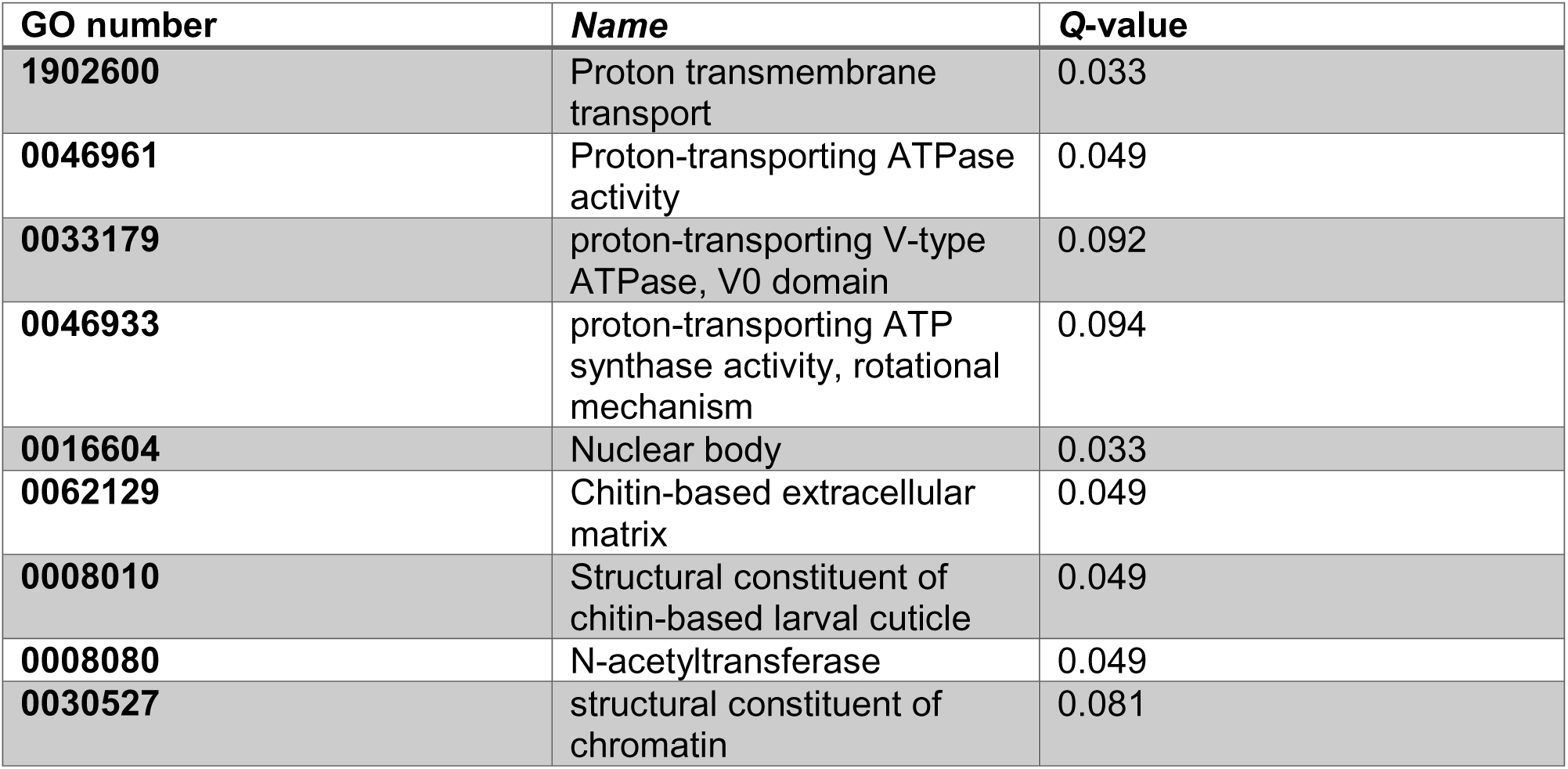
Gene Ontology terms significantly enriched among the top 1% loci in the C_2_ analysis.

**Figure S1.**
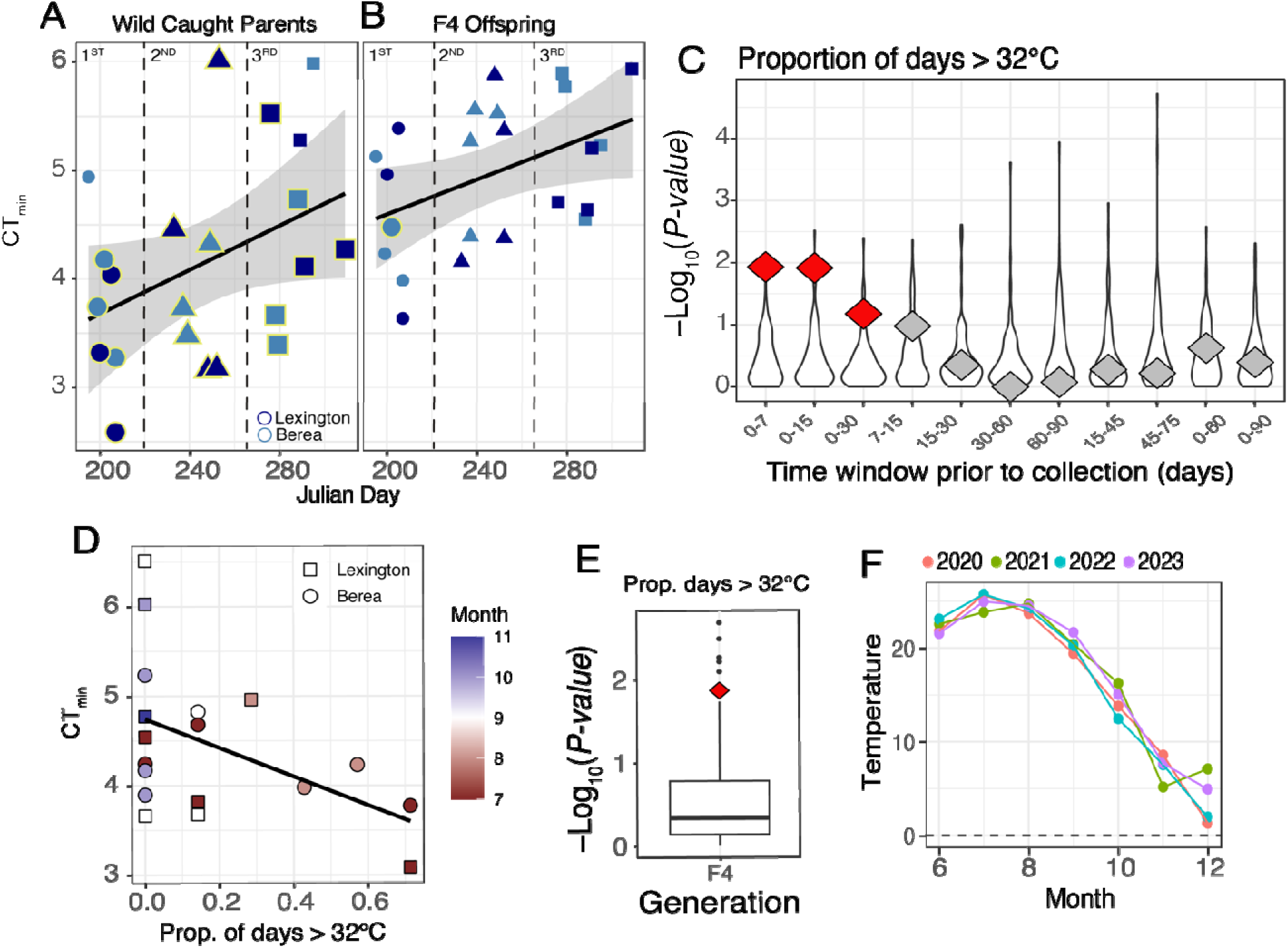
Weather parameters correlated with the trait of CT_min_. (A) CT_min_ values of wild-caught parental flies as a function of Julian date. Colors indicate population of origin. Shapes indicate collection period (circles = first collection, triangles = second collection, squares = third collection). Yellow outlines denote samples that were sequenced for this study. (B) Same as (A), but for F4 offspring reared under common garden conditions. (C) Model search results across environmental time windows for the number of days exceeding 32°C. The y-axis shows model P-values. White violin plots represent the distribution of P-values obtained from permutation tests, whereas diamonds indicate the P-values of the observed data. Red diamonds denote statistically significant models. Best model (0-15 days prior to collection): GLM β_T>32_ = −2.0163, P-value = 0.0145, exceeding 98% of permutations. (D) CT_min_ values of wild-caught parental flies as a function of the proportion of days exceeding 32°C. Shapes indicate collection site. Lower CT_min_ values (greater cold tolerance) were associated with collections made following periods with a higher frequency of days above 32°C. Although seemingly counterintuitive, the most cold tolerant flies were collected early in the season (July-August) and likely represent descendants of individuals that survived overwintering. Consequently, the frequency of days above 32°C serves as a coarse proxy for the seasonal selective history experienced by these populations. (E) Observed and permuted P-values for the 0-15 day window prior to collection in the F4 generation. As with the parenta dataset, the number of days above 32°C during the preceding 0-15 days provided the strongest explanation of variation in CT_min_. generation in the F4: GLM β_T>32_ = −1.0289, P-value = 0.0208, exceeding 95% of permutations. (E) Estimated mean air temperature (2 m above ground level) across years in Lexington, Kentucky, illustrating the consistency of interannual temperature patterns.

**Figure S2.**
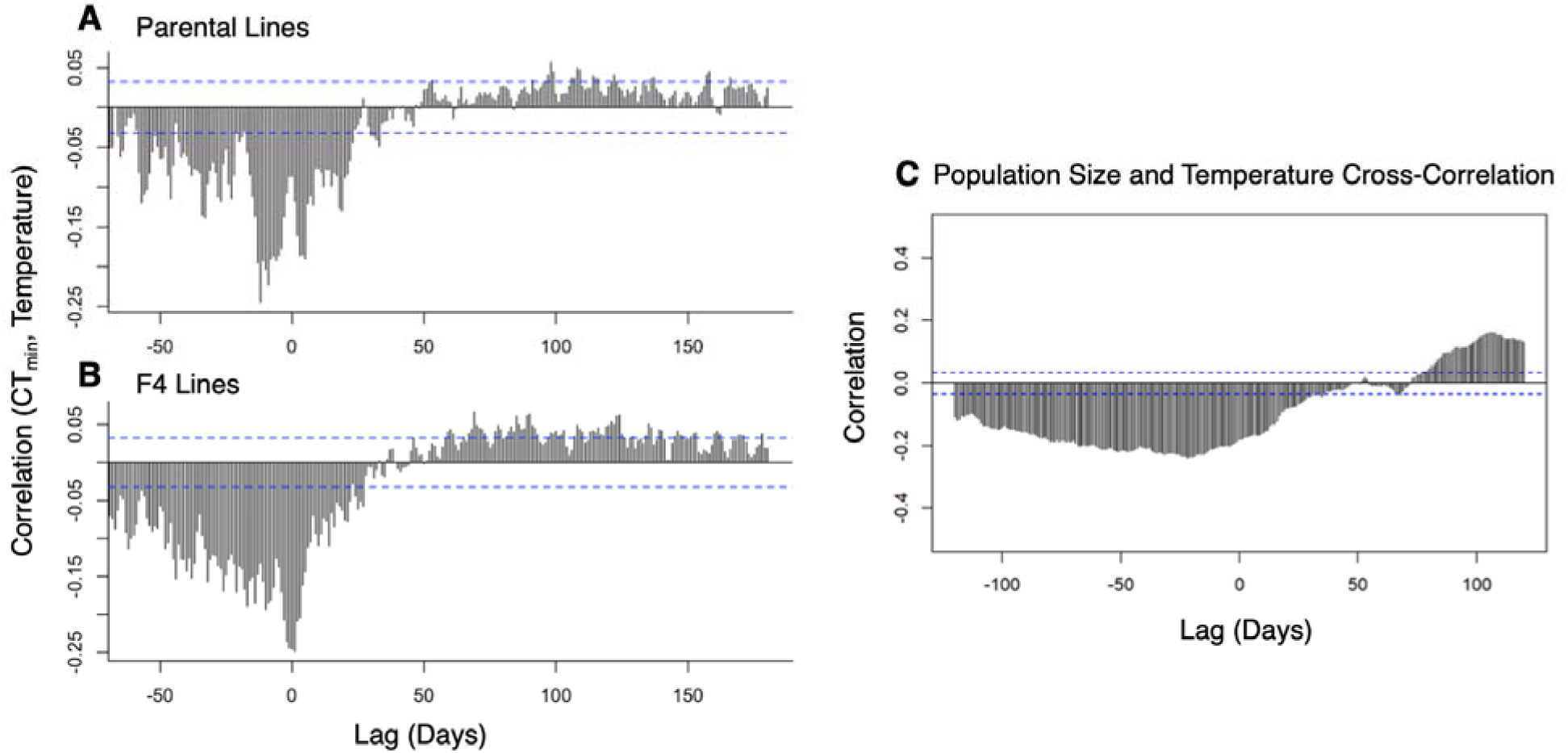
Cross correlation between mean temperature and CT_min._ (A) CT_min_ measured in parental populations, (B) CT_min_ measured in F4 isofemale lines, and (C) D. suzukii population size (from trap data in McCabe et al 2025). Dashed blue lines indicate significance thresholds for the cross-correlation coefficients. Negative correlations indicate that periods with higher temperatures are associated with lower CT_min_ (greater cold tolerance), whereas positive correlations indicate that periods with higher temperatures are associated with higher CT_min_ (reduced cold tolerance). In both parental populations and F4 isofemale lines, the relationship shifts from strongly negative at short lags to positive at lags of approximately 60-120 days. In Panel C, periods of high abundance tend to occur about 60-120 days after periods with higher temperatures, producing positive correlations at positive lags.

**Figure S3.**
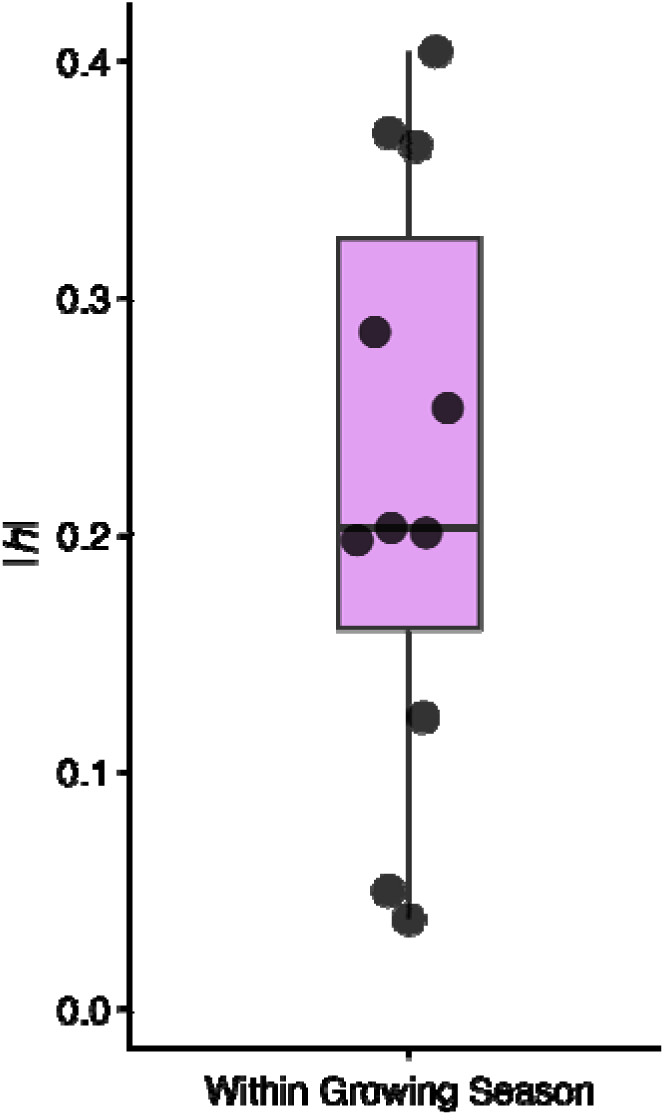
Evolutionary rate analysis for CT_min_. Estimates of evolutionary rates (absolute haldanes, |h|) for wild-caught D. suzukii. Because the number of generations separating successive growing seasons is unknown, evolutionary rates were calculated only for intervals within growing seasons.

**Figure S4.**
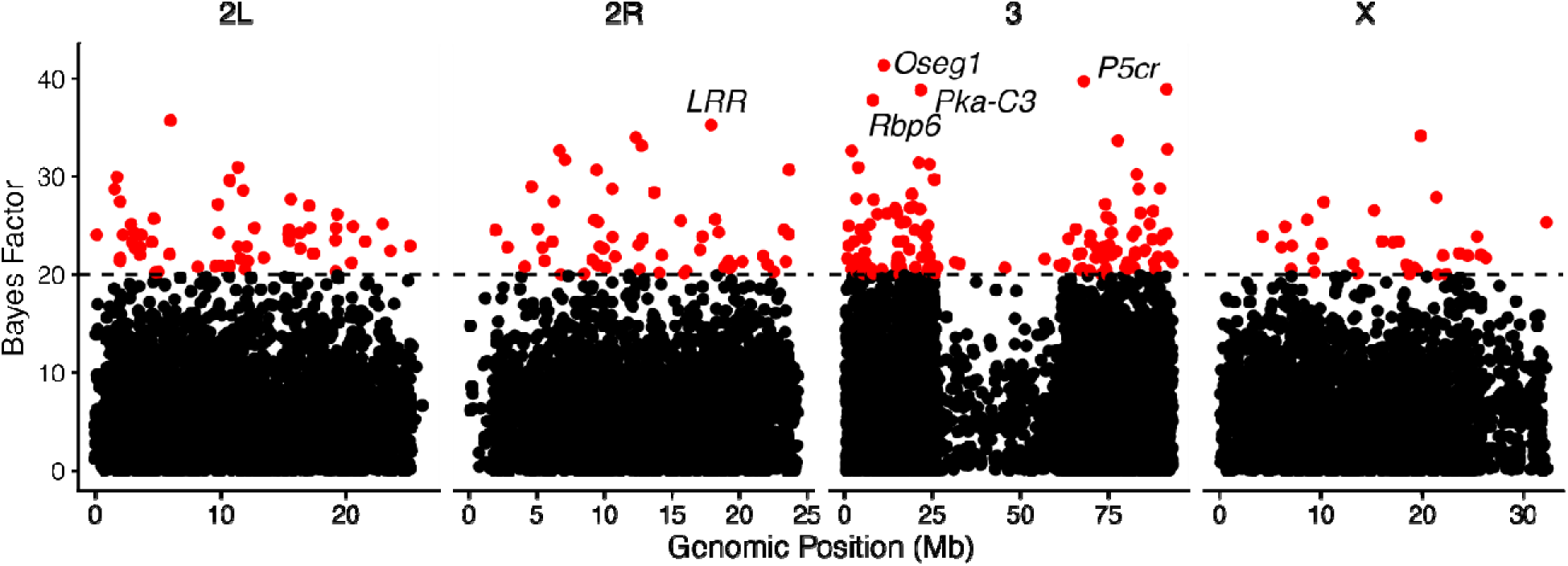
Genetic architecture of CT_min_. Raw association values (i.e., Bayes Factors, BF) results of the pGWAS (i.e., population genome-wide association analysis) using CT_min_ as covariate in BayPass. The y-axis shows the association score measures as BF. The horizontal dashed line represents loci with extremely high evidence for association (BF > 20). A local score analysis of this data does not reveal any genomic regions significant enriched for associated loci, consistent with a highly polygenic genetic architecture.

**Figure S5.**
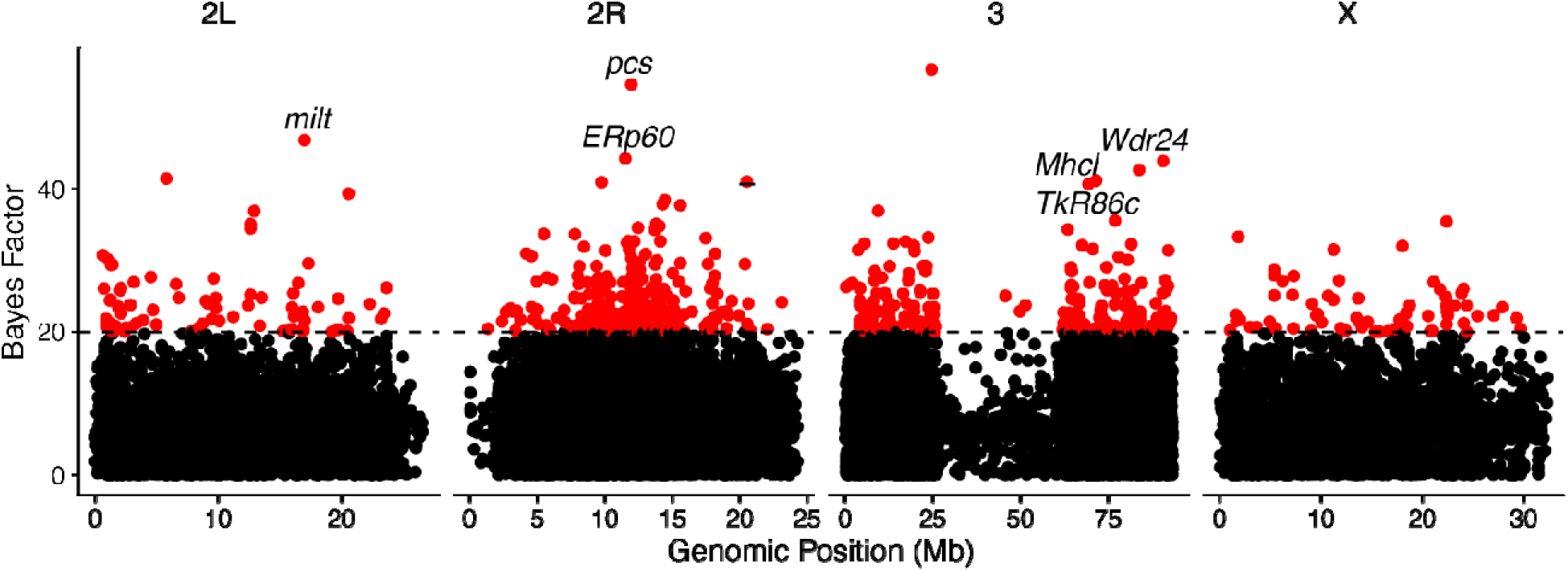
Genome-Environment Association (GEA) analysis of. Raw association values (i.e., Bayes Factors, BF) results of the GEA using as covariate in BayPass. The y-axis shows the association score measures as Bayes factors (BF). The horizontal red line represents loci with extremely high evidence for association (BF > 20). A local score analysis of this data does not reveal any genomic regions significant enriched for associated loci, consistent with a highly polygenic genetic architecture.

## Supplementary Text 1: Phylogeographic Hypotheses Tested

The observed patterns in PCA suggest that D. suzukii populations are characterized by stable, persistent population structures. To test this hypothesis, we conducted model-based demographic inference using the program moments, with a specific focus on the neighboring Virginia (VA) and Kentucky (KY) populations. Given their recent invasion history, our core goal was to determine whether these populations are better described by a panmictic, single-population, model (Fig. TS1-1 A), by a discrete two-population model with migration (symmetrical or asymmetrical; Fig. TS1-1 B, C), or by a model with no migration (Fig. TS1-1 D).

We fitted all demographic models pairwise across every combination of Kentucky and Vermont population pools and calculated the Akaike Information Criterion (AIC) for each model in each comparison. To summarize support across comparisons, we identified the best-fitting model, defined as the model with the lowest AIC, for each pairwise analysis and then quantified the distribution of best-supported models across all comparisons. The results strongly rejected both the panmictic model and the model without migration. Instead, models incorporating gene flow consistently received the greatest support (Fig. TS1-1 E). Among these, the symmetric migration model generally outperformed the asymmetric migration model, although the difference in AIC was modest, indicating only marginally stronger support for symmetric gene flow between populations. These findings support our initial hypothesis that the KY and VA populations are spatially distinct yet connected by ongoing gene flow. Estimated divergence times spanned a broad range, from approximately 31 to 670 years. The upper end of this range is difficult to reconcile with the invasion history of D. suzukii, which was first detected in the continental United States in 2008. In contrast, the lowest divergence time estimates are consistent with the documented timeline of introduction and range expansion. Migration rates were estimated to range from approximately 0.5 to 2.5 effective migrants per generation, indicating continued gene flow among populations despite evidence of demographic subdivision.

**Figure TS1-1:**
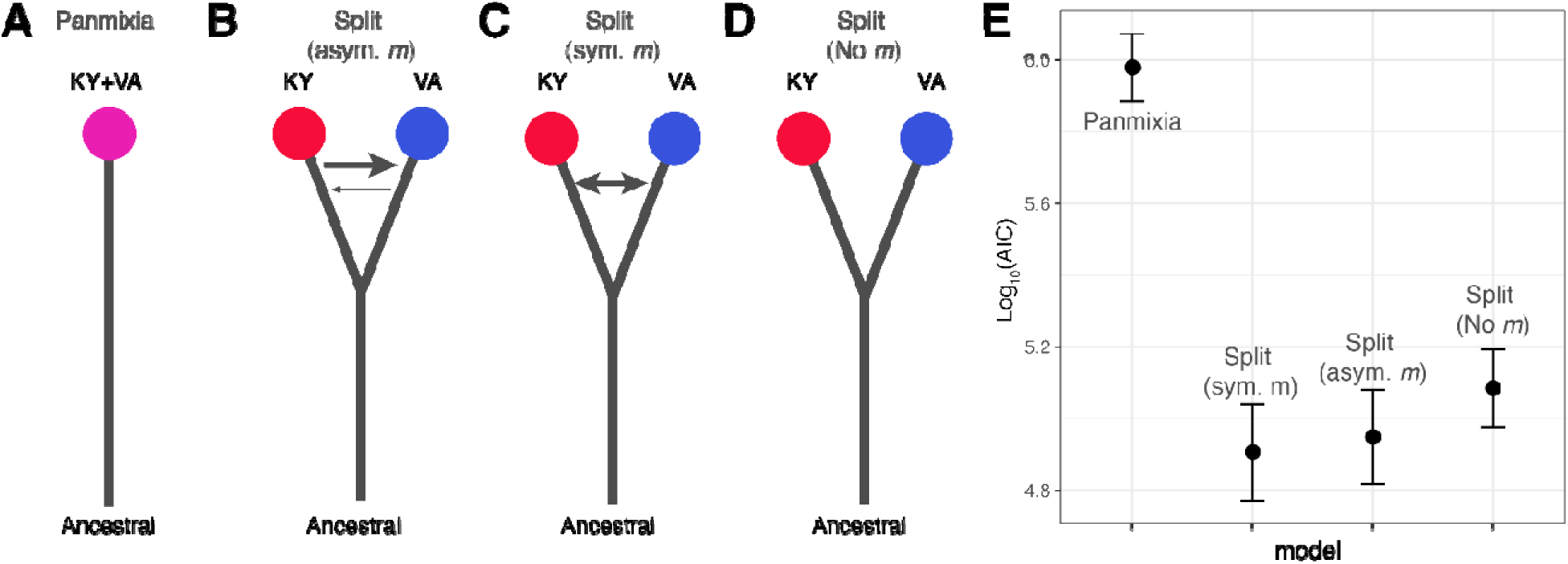
(A) Pictorial representation of a “quasi-panmictic” model for VA and KY. (B) Same as A, but for a discrete two-population model with symmetrical migration. (C) Same as A, but for a discrete two-population model with asymmetrical migration. (D) Same as A, but for a discrete two-population model with no migration. (E) Log_10_(AIC) support for models. Lower AIC is better.

## Supplementary Text 2: Patterns of year-to-year F_ST_ in seasonal simulations

We used SLiM to simulate a population evolving under a boom-and-bust demographic regime. Simulations were conducted in a single deme using a non-Wright-Fisher framework derived from first principles. To model seasonal bottlenecks, we implemented eight winter bottleneck intensities (i.e., “bottleneck param.” in Fig. TS2-1 A) ranging from moderate (∼8% reduction of population size from summer to winter) to extreme (∼99% reduction) and ran 10 replicates per condition.

**Figure TS2-1:**
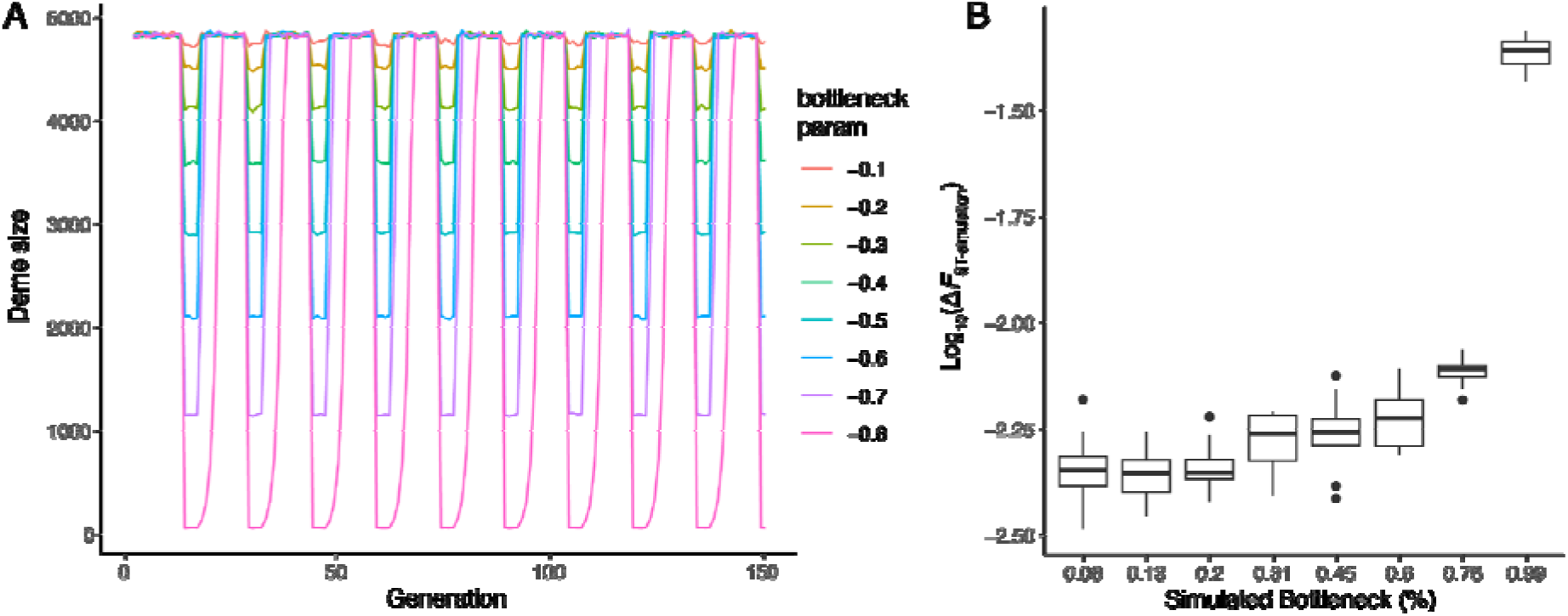
(A) Demographic simulations with boom-and-bust dynamics using various bottleneck parameters. (B) Log_10_ transformed delta values between the within year FST and the across year FST (i.e., overwintering) across different bottleneck magnitudes.

For each simulation, we calculated the difference in F_ST_ between “within-year” (i.e., “across the growing season”) and “between-year” (i.e., “overwintering”) comparisons as a function of bottleneck severity. Across most scenarios, F_ST_ differences were relatively similar, except under the most extreme bottleneck, which produced an order-of-magnitude larger relative divergence (ΔF_ST_ = |F_ST-within y._ - F_ST-between y._ |) between years (Fig. TS2-1 B). This indicates that the sensitivity of F_ST_ to detect demographic contractions declines sharply when bottlenecks are not severe.

In contrast, our empirical data from D. suzukii (see main text) revealed no significant difference between within-year and between-year F_ST_. This lack of signal may reflect a combination of a relatively weak winter bottleneck in D. suzukii and the use of pooled sequencing (Pool-Seq), which can mask subtle allele frequency shifts expected under mild demographic contractions.

Two further caveats are important to note. First, the absolute F_ST_ values in our simulations are entirely theoretical and should not be interpreted as direct estimates for the real data, as they do not incorporate the full complexity of natural demographic and ecological processes. Second, the primary goal of these simulations is to illustrate relative changes in ΔF_ST_ across bottleneck intensities, highlighting the reduced sensitivity of F_ST_ under weaker bottlenecks.

A comprehensive evaluation of how the confluence of Pool-Seq properties, complex demographic histories, and ecological context influence bottleneck detection using F_ST_ is beyond the scope of this study but represents an active area of ongoing research.

## Supplementary Text 3: Differential Expression Analysis

**Figure TS3-1.**
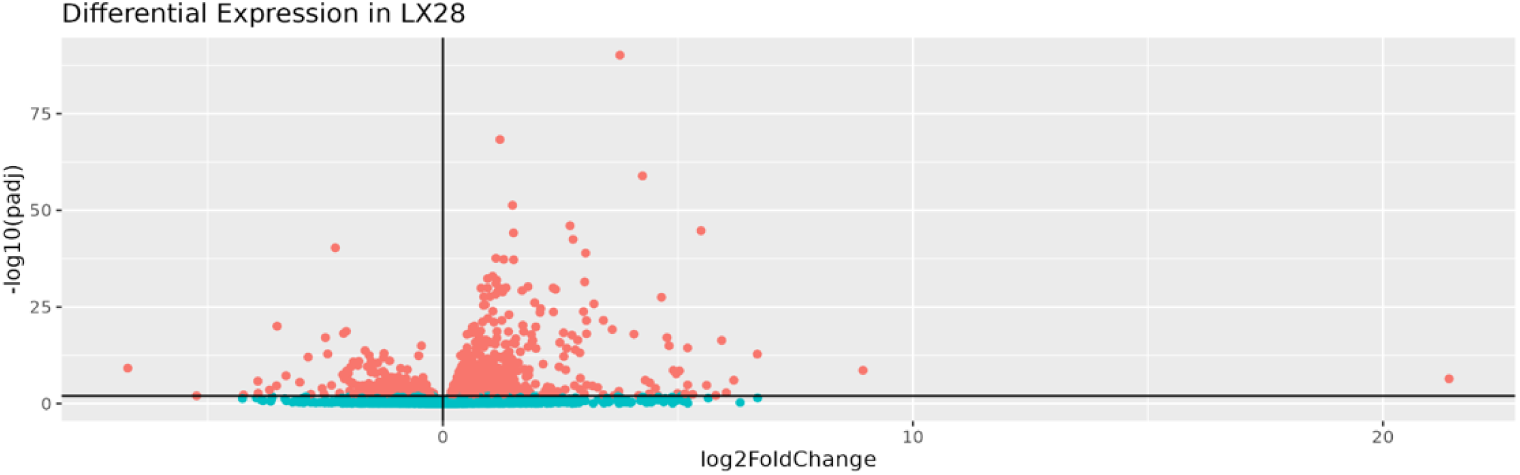
Level of differential expression in LX28 after chilling treatment (red dots signify Padj < 0.01). Dots to the left of the vertical bar represent genes with lower expression levels relative to control. Dots to the right of the vertical bar represent genes with higher expression levels relative to the control. The x-axis uses log2 fold change as a measurement for how highly expressed a gene is. The y-axis uses −log10 of the adjusted p-value as a measurement of how significant the differential expression is.

875 genes show significant differential expression after chilling treatment in a line of D. suzukii (LX28). Flies were collected in Lexington, Kentucky, and subjected to a chilling treatment before RNA-seq (Fig. TS3-1). 602 of these genes are expressed at levels higher than the control and 273 genes are expressed at levels lower than the control. Notable genes include: ImpE2, which is highly expressed (Log_2_[fold change] = 21.41); Ser8, which is expressed at low levels (Log_2_[fold change] = −6.70); and LOC108011169, which has highly significant elevated expression (P = 7.27×10^−91^). ImpE2 codes for ecdysone-inducible gene E2, a gene essential to a complex causing cell death in fruit flies (1-2). Apoptotic genes have previously been shown to be associated with cold stress (3). Ser8 codes for a serine protease, previously found to have lower expression levels in fruit flies selected for chill resistance (4). LOC108011169 codes for an uncharacterized pancreatic triacylglycerol lipase homolog. Triacylglycerol is the main source of metabolic energy in overwintering fruit flies. Triacylglycerols in overwintering fruit flies and flies adapted to colder climates have lower transcription temperatures due to changes in fatty acid composition (5). Additionally, lipid metabolism is well known to be associated with cold stress, due to the need for a more unsaturated phospholipid membrane (6-8).

**Figure TS3-2.**
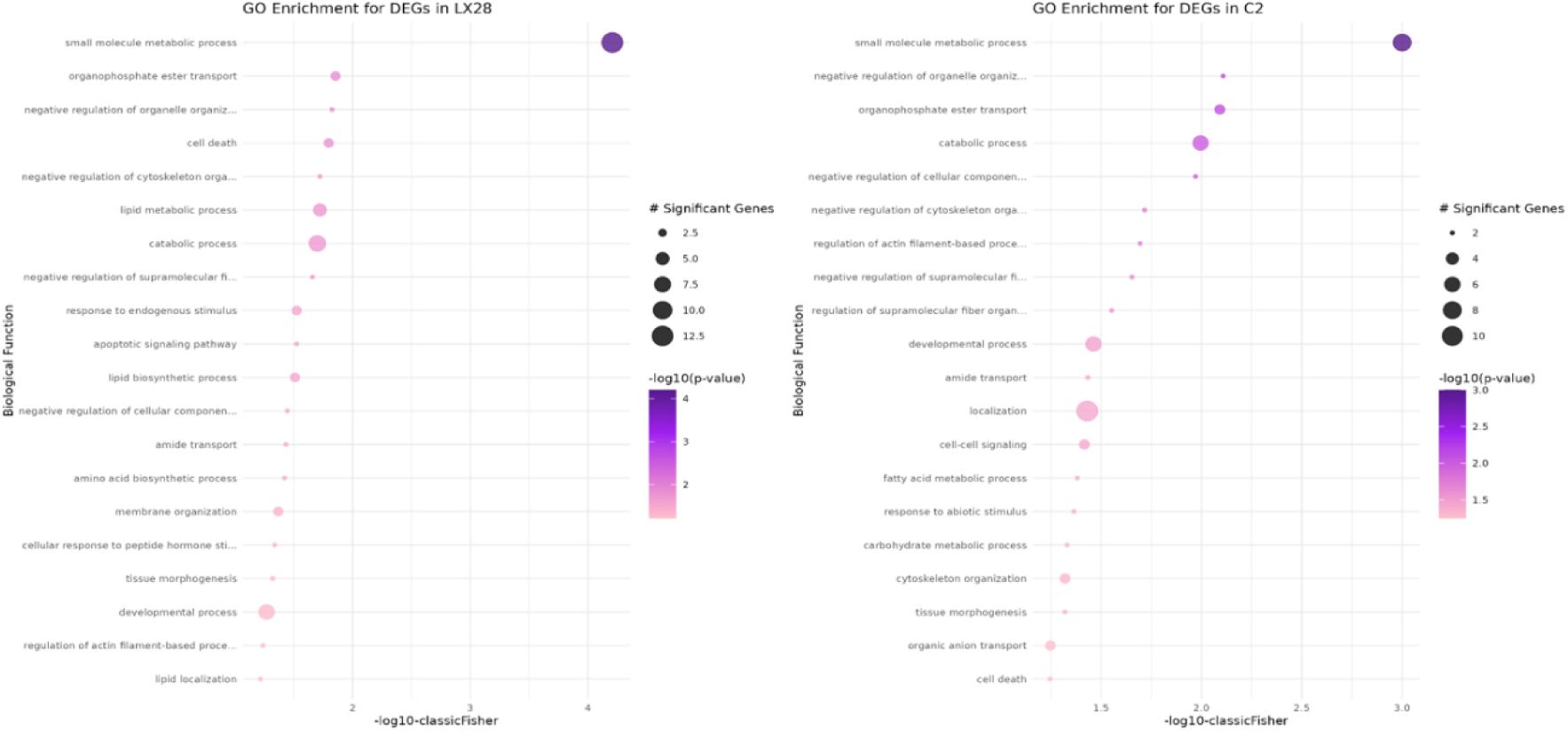
Gene Ontology (GO) enrichment for differentially expressed genes in both LX28 (left) and the C2 dataset (right). Differentially expressed genes were clustered into biological function (y-axis) via Fisher’s exact test. Results are limited to the top 20 GO terms with significant differential expression.

To analyze the relationship between genes differentially expressed in a cold shocked line and genes predicted to be involved in seasonality (Fig. TS3-2), we looked at GO enrichment in the LX28 dataset (left), as well as only genes enriched in both the LX28 and the C_2_ datasets (right). Previous work indicates that the physiological response mechanisms to acute cold stress and long-term cold acclimation are different (9). When comparing the enrichment of genes differentially expressed in both the LX28 dataset and the combined dataset, we find strong similarities. Of note are terms related to cytoskeletal organization and lipid synthesis and localization. As noted above, lipid metabolism is well known to be associated with cold stress. Differential expression in both datasets indicates a possible connection between response to both acute cold stress and longer-term seasonal acclimation to colder temperatures. Long-term exposure to colder temperatures is also known to be associated with changes in cytoskeletal organization in Drosophila (10-11), indicating another possible connection between physiological mechanisms of both acute cold shock response and long-term acclimation response. These results indicate that, while physiological mechanisms between different cold treatments may differ, they may also share similar response pathways.

## References

1. Bellard, C., C. Bernery, and C. Leclerc, Looming extinctions due to invasive species: Irreversible loss of ecological strategy and evolutionary history. Global Change Biology, 2021. 27(20): p. 4967–4979.

2. Marbuah, G., I.-M. Gren, and B. McKie, Economics of harmful invasive species: a review. Diversity, 2014. 6(3): p. 500–523.

3. Pimentel, D., R. Zuniga, and D. Morrison, Update on the environmental and economic costs associated with alien-invasive species in the United States. Ecological economics, 2005. 52(3): p. 273–288.

4. Crystal-Ornelas, R., et al., Economic costs of biological invasions within North America. NeoBiota, 2021. 67: p. 485–510.

5. Jeschke, J.M. and D.L. Strayer, Invasion success of vertebrates in Europe and North America. Proceedings of the National Academy of Sciences, 2005. 102(20): p. 7198–7202.

6. Simberloff, D., The role of propagule pressure in biological invasions. Annual review of ecology, evolution, and systematics, 2009. 40(1): p. 81–102.

7. Bernery, C., et al., Freshwater fish invasions: A comprehensive review. Annual Review of Ecology, Evolution, and Systematics, 2022. 53(1): p. 427–456.

8. Gioria, M., et al., Why are invasive plants successful? Annual Review of Plant Biology, 2023. 74(1): p. 635–670.

9. Zhao, Z., et al., The world’s 100 worst invasive alien insect species differ in their characteristics from related non-invasive species. Journal of Applied Ecology, 2023. 60(9): p. 1929–1938.

10. Sherpa, S. and L. Després, The evolutionary dynamics of biological invasions: A multi-approach perspective. Evolutionary applications, 2021. 14(6): p. 1463–1484.

11. Botero, C.A., et al., Evolutionary tipping points in the capacity to adapt to environmental change. Proceedings of the National Academy of Sciences, 2015. 112(1): p. 184–189.

12. Nunez, J.C., et al., A cosmopolitan inversion facilitates seasonal adaptation in overwintering Drosophila. Genetics, 2024. 226(2): p. iyad207.

13. Bergland, A.O., et al., Genomic evidence of rapid and stable adaptive oscillations over seasonal time scales in Drosophila. PLoS genetics, 2014. 10(11): p. e1004775.

14. Dobzhansky, T., Genetics of natural populations. XXV. Genetic changes in populations of Drosophila pseudoobscura and Drosophila persimilis in some localities in California. Evolution, 1956: p. 82–92.

15. Bogaerts-Márquez, M., et al., Temperature, rainfall and wind variables underlie environmental adaptation in natural populations of Drosophila melanogaster. Molecular ecology, 2021. 30(4): p. 938–954.

16. Machado, H.E., et al., Broad geographic sampling reveals the shared basis and environmental correlates of seasonal adaptation in Drosophila. Elife, 2021. 10: p. e67577.

17. Rudman, S.M., et al., Direct observation of adaptive tracking on ecological time scales in Drosophila. Science, 2022. 375(6586): p. eabj7484.

18. Prileson, E.G., et al., Overwintering drives rapid adaptation in Drosophila with potential costs to insecticide resistance. Evolution, 2025: p. qpaf205.

19. Behrman, E.L., et al., Rapid seasonal evolution in innate immunity of wild Drosophila melanogaster. Proceedings of the Royal Society B: Biological Sciences, 2018. 285(1870): p. 20172599.

20. Pfenninger, M. and Q. Foucault, Population genomic time series data of a natural population suggests adaptive tracking of fluctuating environmental changes. Integrative and Comparative Biology, 2022. 62(6): p. 1812–1826.

21. Behrman, E.L., et al., Seasonal variation in life history traits in two Drosophila species. Journal of Evolutionary Biology, 2015. 28(9): p. 1691–1704.

22. Rajpurohit, S., et al., Adaptive dynamics of cuticular hydrocarbons in Drosophila. Journal of evolutionary biology, 2017. 30(1): p. 66–80.

23. Erickson, P.A., et al., Unique genetic signatures of local adaptation over space and time for diapause, an ecologically relevant complex trait, in Drosophila melanogaster. PLoS Genetics, 2020. 16(11): p. e1009110.

24. Asplen, M.K., et al., Invasion biology of spotted wing Drosophila (Drosophila suzukii): a global perspective and future priorities. Journal of Pest Science, 2015. 88: p. 469–494.

25. Farnsworth, D., et al., Economic analysis of revenue losses and control costs associated with the spotted wing drosophila, Drosophila suzukii (Matsumura), in the California raspberry industry. Pest management science, 2017. 73(6): p. 1083–1090.

26. McCabe, E.A., et al., Environmental conditions influencing seasonal population dynamics of Drosophila suzukii (Diptera: Drosophilidae) in mid-latitude organic farms. Agricultural and Forest Entomology, 2025.

27. Lewald, K.M., et al., Population genomics of Drosophila suzukii reveal longitudinal population structure and signals of migrations in and out of the continental United States. G3, 2021. 11(12): p. jkab343.

28. Camus, L., et al., Adaptive Challenges of Past and Future Invasion of Drosophila suzukii: Insights From Novel Genomic Resources and Statistical Methods Combining Individual and Pool Sequencing Data. Molecular Ecology, 2025: p. e70192.

29. Olazcuaga, L., et al., A whole-genome scan for association with invasion success in the fruit fly Drosophila suzukii using contrasts of allele frequencies corrected for population structure. Molecular biology and evolution, 2020. 37(8): p. 2369–2385.

30. Adrion, J.R., et al., Drosophila suzukii: the genetic footprint of a recent, worldwide invasion. Molecular biology and evolution, 2014. 31(12): p. 3148–3163.

31. Gautier, M., et al., f-Statistics estimation and admixture graph construction with Pool-Seq or allele count data using the R package poolfstat. Molecular Ecology Resources, 2022. 22(4): p. 1394–1416.

32. Fraimout, A., et al., Deciphering the routes of invasion of Drosophila suzukii by means of ABC random forest. Molecular biology and evolution, 2017. 34(4): p. 980–996.

33. Nelson, T.D., et al., Spotted-wing drosophila, Drosophila suzukii (Diptera: Drosophilidae), exhibits large-scale spatial genetic structure across Canada but not fruit host–associated genetic structure. The Canadian Entomologist, 2023. 155: p. e37.

34. Schmidt, P.S. and D.R. Conde, Environmental heterogeneity and the maintenance of genetic variation for reproductive diapause in Drosophila melanogaster. Evolution, 2006. 60(8): p. 1602–1611.

35. Min, J., et al., A general model for the evolution of thermal performance curves with application to real time-series data. bioRxiv, 2026: p. 2026.06. 17.733045.

36. Sinclair, B.J., C.M. Williams, and J.S. Terblanche, Variation in thermal performance among insect populations. Physiological and biochemical zoology, 2012. 85(6): p. 594–606.

37. Ayrinhac, A., et al., Cold adaptation in geographical populations of Drosophila melanogaster: phenotypic plasticity is more important than genetic variability. Functional Ecology, 2004: p. 700–706.

38. Kinnison, M.T. and A.P. Hendry, The pace of modern life II: from rates of contemporary microevolution to pattern and process. Genetica, 2001. 112(1): p. 145–164.

39. Marshall, K.E. and B.J. Sinclair, Repeated stress exposure results in a survival– reproduction trade-off in Drosophila melanogaster. Proceedings of the Royal Society B: Biological Sciences, 2009. 277(1683): p. 963.

40. Schou, M.F., et al., Evolutionary trade-offs between heat and cold tolerance limit responses to fluctuating climates. Science Advances, 2022. 8(21): p. eabn9580.

41. Lee, G., et al., A large-effect fitness trade-off across environments is explained by a single mutation affecting cold acclimation. Proceedings of the National Academy of Sciences, 2024. 121(6): p. e2317461121.

42. Ørsted, M., et al., Global analysis of the seasonal abundance of the invasive pest Drosophila suzukii reveal temperature extremes determine population activity potential. Pest management science, 2021. 77(10): p. 4555–4563.

43. Stacconi, V.R., Drosophila suzukii (spotted wing drosophila). 2022.

44. McCabe, E.A., L.N. Unfried, and N.M. Teets, Survival and nutritional requirements for overwintering Drosophila suzukii (Diptera: Drosophilidae) in Kentucky. Environmental Entomology, 2023. 52(6): p. 1071–1081.

45. Ueno, T., et al., Rapid seasonal changes in phenotypes in a wild Drosophila population. Scientific Reports, 2023. 13(1): p. 21940.

46. Lenormand, T., Gene flow and the limits to natural selection. Trends in ecology & evolution, 2002. 17(4): p. 183–189.

47. Fariello, M.I., et al., Accounting for linkage disequilibrium in genome scans for selection without individual genotypes: the local score approach. Molecular ecology, 2017. 26(14): p. 3700–3714.

48. Daborn, P., et al., A single P450 allele associated with insecticide resistance in Drosophila. Science, 2002. 297(5590): p. 2253–2256.

49. Le Goff, G. and F. Hilliou, Resistance evolution in Drosophila: the case of CYP6G1. Pest Management Science, 2017. 73(3): p. 493–499.

50. Gul, H., et al., Fitness costs of resistance to insecticides in insects. Frontiers in physiology, 2023. 14: p. 1238111.

51. Zhang, L.J., et al., Trade-off between thermal tolerance and insecticide resistance in Plutella xylostella. Ecology and Evolution, 2015. 5(2): p. 515–530.

52. Scheuermann, E.A. and D.P. Smith, Odor-specific deactivation defects in a Drosophila odorant-binding protein mutant. Genetics, 2019. 213(3): p. 897–909.

53. Park, K., et al., Molecular and cellular organization of odorant binding protein genes in Drosophila. Heliyon, 2024. 10(9).

54. Olazcuaga, L., et al., Rapid and transient evolution of local adaptation to seasonal host fruits in an invasive pest fly. Evolution Letters, 2022. 6(6): p. 490–505.

55. Takeuchi, K.-i., et al., Changes in temperature preferences and energy homeostasis in dystroglycan mutants. Science, 2009. 323(5922): p. 1740–1743.

56. Thanh, M.T., et al., Drosophila model for studying the link between lipid metabolism and development. Frontiers in Bioscience-Landmark, 2020. 25(1): p. 147–158.

57. Adachi, Y., et al., AGPAT2 acts at the crossroads of lipid biosynthesis and DRP1-mediated ER morphogenesis. Nature communications, 2025. 16(1): p. 11473.

58. Vial, T., et al., Dengue virus reduces AGPAT1 expression to alter phospholipids and enhance infection in Aedes aegypti. PLoS Pathogens, 2019. 15(12): p. e1008199.

59. Enriquez, T. and N.M. Teets, Lipid properties and metabolism in response to cold. 2025.

60. Rider, M.H., et al., AMP-activated protein kinase and metabolic regulation in cold-hardy insects. Journal of insect physiology, 2011. 57(11): p. 1453–1462.

61. Johnson, E.C., et al., Altered metabolism and persistent starvation behaviors caused by reduced AMPK function in Drosophila. PloS one, 2010. 5(9): p. e12799.

62. Colautti, R.I. and S.C. Barrett, Rapid adaptation to climate facilitates range expansion of an invasive plant. science, 2013. 342(6156): p. 364–366.

63. van Boheemen, L.A., D.Z. Atwater, and K.A. Hodgins, Rapid and repeated local adaptation to climate in an invasive plant. New Phytologist, 2019. 222(1): p. 614–627.

64. Martin, R.A., et al., When will a changing climate outpace adaptive evolution? Wiley Interdisciplinary Reviews: Climate Change, 2023. 14(6): p. e852.

65. Qin, W., et al., Hugin-AstA circuitry is a novel central energy sensor that directly regulates sweet sensation in Drosophila and mouse. Elife, 2026. 14: p. RP108551.

66. Lecheta, M.C., et al., Integrating GWAS and transcriptomics to identify the molecular underpinnings of thermal stress responses in Drosophila melanogaster. Frontiers in genetics, 2020. 11: p. 658.

67. Sparks, A.H., nasapower: a NASA POWER global meteorology, surface solar energy and climatology data client for R. Journal of Open Source Software, 2018. 3(30): p. 1035.

68. Trapletti, A. and K. Hornik, Time series analysis and computational finance (R Package version 0.54). 2023.

69. Gingerich, P.D., Quantification and comparison of evolutionary rates. American Journal of Science, 1993. 293(A): p. 453–478.

70. Gautier, M., Efficient k-mer based curation of raw sequence data: application in Drosophila suzukii. Peer Community Journal, 2023. 3.

71. Vasimuddin, M., et al. Efficient architecture-aware acceleration of BWA-MEM for multicore systems. in 2019 IEEE international parallel and distributed processing symposium (IPDPS). 2019. IEEE.

72. Danecek, P., et al., Twelve years of SAMtools and BCFtools. Gigascience, 2021. 10(2): p. giab008.

73. Camus, L., et al., New insights into past and future invasion of Drosophila suzukii using novel genomic resources and statistical methods to combine individual and pool sequencing data. bioRxiv, 2024: p. 2024.10. 11.617812.

74. Garrison, E. and G. Marth, Haplotype-based variant detection from short-read sequencing. arXiv preprint arXiv:1207.3907, 2012.

75. Cingolani, P., et al., A program for annotating and predicting the effects of single nucleotide polymorphisms, SnpEff: SNPs in the genome of Drosophila melanogaster strain w1118; iso-2; iso-3. fly, 2012. 6(2): p. 80–92.

76. Gupta, S., et al., Quantifying similarity between motifs. Genome biology, 2007. 8(2): p. R24.

77. Hivert, V., et al., Measuring genetic differentiation from pool-seq data. Genetics, 2018. 210(1): p. 315–330.

78. Gautier, M., M. Coronado-Zamora, and R. Vitalis, Estimating hierarchical F–statistics from Pool–Seq data. bioRxiv, 2024: p. 2024.11.22.624688.

79. Jouganous, J., et al., Inferring the joint demographic history of multiple populations: beyond the diffusion approximation. Genetics, 2017. 206(3): p. 1549–1567.

80. Kapun, M., et al., Drosophila evolution over space and time (DEST): a new population genomics resource. Molecular biology and evolution, 2021. 38(12): p. 5782–5805.

81. Nunez, J.C., et al., Footprints of worldwide adaptation in structured populations of Drosophila melanogaster through the expanded dest 2.0 genomic resource. Molecular biology and evolution, 2025. 42(8): p. msaf132.

82. Gautier, M., Genome-wide scan for adaptive divergence and association with population-specific covariates. Genetics, 2015. 201(4): p. 1555–1579.

83. Andrews, S., FastQC: a quality control tool for high throughput sequence data. 2010. 2017.

84. Bolger, A.M., M. Lohse, and B. Usadel, Trimmomatic: a flexible trimmer for Illumina sequence data. Bioinformatics, 2014. 30(15): p. 2114–2120.

85. Kim, D., et al., Graph-based genome alignment and genotyping with HISAT2 and HISAT-genotype. Nature biotechnology, 2019. 37(8): p. 907–915.

86. Love, M.I., W. Huber, and S. Anders, Moderated estimation of fold change and dispersion for RNA-seq data with DESeq2. Genome biology, 2014. 15(12): p. 550.

87. Alexa, A. and J. Rahnenfuhrer, topGO: Enrichment analysis for Gene Ontology. R package version 2.28. 0. Cranio, 2016. 2: p. 289–300.

## References (Supplementary Methods)

1. McCabe, E.A., L.N. Unfried, and N.M. Teets, Survival and nutritional requirements for overwintering Drosophila suzukii (Diptera: Drosophilidae) in Kentucky. Environmental Entomology, 2023. 52(6): p. 1071–1081.

2. McCabe, E.A., et al., Environmental conditions influencing seasonal population dynamics of Drosophila suzukii (Diptera: Drosophilidae) in mid latitude organic farms. Agricultural and Forest Entomology, 2025.

3. Lecheta, M.C., et al., Integrating GWAS and transcriptomics to identify the molecular underpinnings of thermal stress responses in Drosophila melanogaster. Frontiers in genetics, 2020. 11: p. 658.

4. Huey, R., et al., A method for rapid measurement of heat or cold resistance of small insects. Functional Ecology, 1992: p. 489–494.

5. Awde, D.N., et al., High-throughput assays of critical thermal limits in insects. Journal of Visualized Experiments, 2020(160).

6. Bergland, A.O., et al., Genomic evidence of rapid and stable adaptive oscillations over seasonal time scales in Drosophila. PLoS genetics, 2014. 10(11): p. e1004775.

7. Nunez, J.C., et al., A cosmopolitan inversion facilitates seasonal adaptation in overwintering Drosophila. Genetics, 2024. 226(2): p. iyad207.

8. Machado, H.E., et al., Broad geographic sampling reveals the shared basis and environmental correlates of seasonal adaptation in Drosophila. Elife, 2021. 10: p. e67577.

9. Sparks, A.H., nasapower: a NASA POWER global meteorology, surface solar energy and climatology data client for R. Journal of Open Source Software, 2018. 3(30): p. 1035.

10. Trapletti, A. and K. Hornik, Time series analysis and computational finance (R Package version 0.54). 2023.

11. Camus, L., et al., Adaptive Challenges of Past and Future Invasion of Drosophila suzukii: Insights From Novel Genomic Resources and Statistical Methods Combining Individual and Pool Sequencing Data. Molecular Ecology, 2025: p. e70192.

12. Gautier, M., Efficient k-mer based curation of raw sequence data: application in Drosophila suzukii. Peer Community Journal, 2023. 3.

13. Vasimuddin, M., et al. Efficient architecture-aware acceleration of BWA-MEM for multicore systems. in 2019 IEEE international parallel and distributed processing symposium (IPDPS). 2019. IEEE.

14. Danecek, P., et al., Twelve years of SAMtools and BCFtools. Gigascience, 2021. 10(2): p. giab008.

15. Sario, S., J. Melo-Ferreira, and C. Santos, Winter Is (Not) Coming: Is Climate Change Helping Drosophila suzukii Overwintering? Biology, 2023. 12(7): p. 907.

16. Olazcuaga, L., et al., A whole-genome scan for association with invasion success in the fruit fly Drosophila suzukii using contrasts of allele frequencies corrected for population structure. Molecular biology and evolution, 2020. 37(8): p. 2369–2385.

17. Garrison, E. and G. Marth, Haplotype-based variant detection from short-read sequencing. arXiv preprint arXiv:1207.3907, 2012.

18. Gautier, M., et al., f Statistics estimation and admixture graph construction with Pool Seq or allele count data using the R package poolfstat. Molecular Ecology Resources, 2022. 22(4): p. 1394–1416.

19. Cingolani, P., et al., A program for annotating and predicting the effects of single nucleotide polymorphisms, SnpEff: SNPs in the genome of Drosophila melanogaster strain w1118; iso-2; iso-3. fly, 2012. 6(2): p. 80–92.

20. Gupta, S., et al., Quantifying similarity between motifs. Genome biology, 2007. 8(2): p. R24.

21. Hivert, V., et al., Measuring genetic differentiation from pool-seq data. Genetics, 2018. 210(1): p. 315–330.

22. Gautier, M., M. Coronado-Zamora, and R. Vitalis, Estimating hierarchical F–statistics from Pool–Seq data. bioRxiv, 2024: p. 2024.11.22.624688.

23. Jouganous, J., et al., Inferring the joint demographic history of multiple populations: beyond the diffusion approximation. Genetics, 2017. 206(3): p. 1549–1567.

24. Kapun, M., et al., Drosophila evolution over space and time (DEST): a new population genomics resource. Molecular biology and evolution, 2021. 38(12): p. 5782–5805.

25. Nunez, J.C., et al., Footprints of worldwide adaptation in structured populations of Drosophila melanogaster through the expanded dest 2.0 genomic resource. Molecular biology and evolution, 2025. 42(8): p. msaf132.

26. Kolaczkowski, B., et al., Genomic differentiation between temperate and tropical Australian populations of Drosophila melanogaster. Genetics, 2011. 187(1): p. 245–260.

27. Feder, A.F., D.A. Petrov, and A.O. Bergland, LDx: estimation of linkage disequilibrium from high-throughput pooled resequencing data. PloS one, 2012. 7(11): p. e48588.

28. Gautier, M., et al., Estimation of population allele frequencies from next generation sequencing data: pool versus individual based genotyping. Molecular Ecology, 2013. 22(14): p. 3766–3779.

29. Haller, B.C. and P.W. Messer, SLiM 4: multispecies eco-evolutionary modeling. The American Naturalist, 2023. 201(5): p. E127–E139.

30. Baumdicker, F., et al., Efficient ancestry and mutation simulation with msprime 1.0. Genetics, 2022. 220(3): p. iyab229.

31. Norberg, J., Biodiversity and ecosystem functioning: a complex adaptive systems approach. Limnology and Oceanography, 2004. 49(4part2): p. 1269–1277.

32. Gautier, M., Genome-wide scan for adaptive divergence and association with population-specific covariates. Genetics, 2015. 201(4): p. 1555–1579.

33. Andrews, S., FastQC: a quality control tool for high throughput sequence data. 2010. 2017.

34. Ewels, P., et al., MultiQC: summarize analysis results for multiple tools and samples in a single report. Bioinformatics, 2016. 32(19): p. 3047–3048.

35. Bolger, A.M., M. Lohse, and B. Usadel, Trimmomatic: a flexible trimmer for Illumina sequence data. Bioinformatics, 2014. 30(15): p. 2114–2120.

36. Kim, D., et al., Graph-based genome alignment and genotyping with HISAT2 and HISAT-genotype. Nature biotechnology, 2019. 37(8): p. 907–915.

37. Li, H., et al., The sequence alignment/map format and SAMtools. bioinformatics, 2009. 25(16): p. 2078–2079.

38. Okonechnikov, K., A. Conesa, and F. García-Alcalde, Qualimap 2: advanced multi-sample quality control for high-throughput sequencing data. Bioinformatics, 2016. 32(2): p. 292–294.

39. Liao, Y., G.K. Smyth, and W. Shi, featureCounts: an efficient general purpose program for assigning sequence reads to genomic features. Bioinformatics, 2014. 30(7): p. 923–930.

40. Love, M.I., W. Huber, and S. Anders, Moderated estimation of fold change and dispersion for RNA-seq data with DESeq2. Genome biology, 2014. 15(12): p. 550.

41. Alexa, A. and J. Rahnenfuhrer, topGO: Enrichment analysis for Gene Ontology. R package version 2.28. 0. Cranio, 2016. 2: p. 289–300.

## References (Supplementary Text 3)

1. S. Ahmad, I.M. Jami, C.C. Jaworski, Y. Luo, Comparative transcriptomics of the irradiated melon fly (Zeugodacus cucurbitae) reveal key developmental genes. Front. Physiol. 14, (2023).

2. V. Yin, C. Thummel. Mechanisms of steroid-triggered programmed cell death in Drosophila. Semin. Cell Dev. Biol. 16, 237–243 (2005).

3. F. Saeidi, H. Colinet, Thermal stress-induced cell death pathways in Drosophila: molecular insights into apoptosis, necrosis, and autophagy. Insect Biochemistry and Molecular Biology 194, (2026).

4. M. Telonis-Scott, R. Hallas, S.W. McKechnie, C.W. Wee, A.A. Hoffmann, Selection for cold resistance alters gene transcript levels in Drosophila melanogaster. J Insect Physiol 55, 549–555 (2009).

5. T. Ohtsu, C. Katagiri, M.T. Kimura, S.H. Hori, Cold adaptations in Drosophila: Qualitative changes of triacylglycerols with relation to overwintering. Journal of Biological Chemistry 268, 1830–1834 (1993).

6. S.A.L. Hayward, B. Manso, A.R. Cossins, Molecular basis of chill resistance adaptations in poikilothermic animals. J Exp Biol 217, 6–15 (2014).

7. J. Overgaard, J.G. Sørensen, S.O. Petersen, V. Loeschcke, M. Holmstrup, Reorganization of membrane lipids during fast and slow cold hardening in Drosophila melanogaster. Physiological Entomology 31, 328–335 (2006).

8. J. Overgaard, J.G. Sørensen, S.O. Petersen, V. Loeschcke, M. Holmstrup, Changes in membrane lipid composition following rapid cold hardening in Drosophila melanogaster. J Insect Physiol 51, 1173–1182 (2005).

9. N. M. Teets, D. L. Denlinger, Physiological mechanisms of seasonal and rapid cold-hardening in insects. Physiological Entomology 38, 105–116 (2013).

10. H. A. MacMillan, et al., Cold acclimation wholly reorganizes the Drosophila melanogaster transcriptome and metabolome. Sci Rep 6, 28999 (2016).

11. D. J. Parker, et al., How consistent are the transcriptome changes associated with cold acclimation in two species of the Drosophila virilis group? Heredity (Edinb) 115, 13–21 (2015).

